# Emodin modulates PI3K-AKT pathway to inhibit proliferation, invasion and induce apoptosis in glioma cells

**DOI:** 10.1101/2024.02.23.580874

**Authors:** Ashaq Hussain Mir, Mujeeb Zafar Banday, Fayeem Aadil, Shabir Ahmad Ganie, Ehtishamul Haq

## Abstract

Glioma is a type of tumor that begins in glial cells and occurs in the brain and spinal cord. Glioma forms a major health challenge worldwide. They are hard to treat, not only because of the deregulation in multiple signaling transduction pathways affecting various cellular processes but also because they are not contained in a well-defined mass with clear borders. One of the main pathways deregulated in glioma is PI3K-AKT and its associated downstream targets like NF-ĸB which affects different proteins/transcription factors influencing many aspects of gliomagenesis like epithelial to mesenchymal transition (EMT). A combination of *in-silico* and *in-vitro* approaches targeted against specific catalytic isoform (p110δ) of Class IA PI3K with potent and selective inhibitors would maximize the chances of tumor regression. We adopted an in-silico approach to screen a range of natural molecules for a potent p110δ inhibitor and among them, “emodin” was found to be a potential candidate. In vitro, emodin treatment inhibits proliferation, induces apoptosis, modulates astrocytic phenotype, and decreases cell density of glioma cells. Emodin induces changes in the astrocytic phenotype of glioma cells to elongated form with rounded-off, shrunken-down morphology. Emodin was found to contribute to ROS production which leads to apoptosis of glioma cells. The apoptosis induced by emodin was confirmed by propidium iodide staining and ascertained by FACS analysis. We evaluated the effect of emodin on various proteins of PI3K-AKT and downstream targets. We found that emodin treatment decreases the expression of p-AKT, increases expression levels of Iĸ-B, inhibits nuclear translocation of NF-ĸB, and upregulates the phosphorylated form of GSK-3β. Changes at the molecular level of these proteins result in the inhibition/degradation of downstream proteins and transcription factors associated with the growth and proliferation of glioma cells. Inhibition of nuclear translocation of NF-ĸB also inhibits nuclear activation of various protumorigenic signaling pathway mediators involved in EMT such as N-cadherin, β-catenin, Claudin-1. These EMT markers promote invasion, proliferation, migration, and growth in glioma cells. Emodin treatment resulted in changed expression profiles of these EMT markers involved in promoting gliomagenesis. In essence these results suggest that in-vitro emodin treatment remarkably reduces the proliferation of glioma cells possibly targeting multiple pathways involved in tumor growth, proliferation, and development, supporting the rationale and relevance of using multipronged strategies for effective treatment of glioma.

## 1. Introduction

Brain cancer and other cancers of the nervous system remain the 10^th^ leading cause of death for both men and women in the United States only[1]. The average annual incidence (age adjusted) rate of malignant brain and other CNS tumors was found to be 7.08 per 100,000 in 2012-2016. Gliomas are the most common primary tumors and account for about 26% of all CNS tumors in adolescents and young adults and approximately 45.7% of tumors in the age group of 0-19 years in the United States [2, 3]. According to the CBTRUS report, the five-year survival rate for malignant CNS tumors and glioblastoma is 35.8% and 6.8% respectively in the United States[3].

Gliomas are a broad range of primary tumors that appear in the brain or spinal cord from glial cells. These tumors are neuroectodermal in origin [4]. They are thought to originate from stem cells or glial progenitor cell which gradually develops glial features or gliogenic cancer stem cells upon neoplastic transformation due to accumulation and/or incorporation of mutation[5, 6]. Gliomas are characterized by elevated proliferative potential, infiltrative growth tendency, intratumoral heterogeneity, resistance to apoptosis, genomic instability, and immense tumor recurrence [7, 8] and thus remain practically incurable. The standard regimen for treating gliomas includes maximal surgical removal, preceded by radiation therapy with temozolomide (TMZ), an orally prescribed alkylating chemotherapeutic agent, and then adjuvant TMZ chemotherapy (National Comprehensive Cancer Network [NCCN], 2015).The most aggressive, most common, and most frequent form of malignant tumor of the brain is glioblastoma, commonly known as glioblastoma multiforme (GBM). Accordingly, WHO has classified GBM as a grade IV tumor [9]. Macroscopical observation of GBM reveals heterogeneity featuring necrosis, multifocal hemorrhage, and gelatinous and cystic areas [10, 11].

In glioma cells, the over-active PI3K/Akt pathway leads to tumor progression, rapid growth, and multidrug resistance. Inhibiting the PI3k pathway at different nodal points with inhibitors individually or in combination results in reduced tumor progression and glioma cell death. The PI3K family is composed of four classes according to their sequence homology and substrate preference [12, 13]. All class IA PI3Ks are heterodimeric molecules composed of an 85 kD regulatory subunit (p85) and a 110 kD catalytic subunit (p110α, p110β, and p110δ).

Interestingly, p110δ is highly expressed in cancer cell lines, including breast, lung, and neuroblastoma-derived cells. p110δ has been reported to be a key factor for glioma cell migration and invasion and knockdown strategies of p110δ have proved to reduce the migration and invasion capacity of certain glioma cells [14]. It has also been reported that the p110δ PI3K isoform inhibits the tumor suppressor activity of PTEN through a negative signaling cascade that involves RhoA /ROCK inhibition. Besides, p110δ activation positively regulates p190RhoGAP activity, which in turn results in the buildup of p27 in the cytoplasm [15]. Once PI3K is phosphorylated, it leads to the activation of AKT/Protein kinase B which is the nodal molecule for various signaling pathways. AKT leads to the activation of various signaling molecules with varying effector functions. One of the effector molecules is NF-ĸB. NF-ĸB signaling pathway encompasses several mechanisms that are activated by a range of stimuli and ultimately congregate on transcription factors belonging to the NF-ĸB family. NF-ĸB dimer complexes are expressed in a variety of cell types and are kept in an inactive state (non-DNA bound state) in the cytoplasm by binding to specific inhibitors of NF-ĸB (Iĸ-B), till activated by corresponding activating signal. In GBM, there is aberrant and constitutive activation of NF-ĸB [16–20] and several mechanisms have been offered to explain it. For example, RTK particularly PDGFR and EGFR, reported to be regularly and aberrantly activated in glioblastoma have been associated with NF-ĸB activation. NF-ĸB signaling pathway has been implicated in promoting as well as maintaining EMT in cancerous and healthy cells. This function is carried out by regulating the expression of genes implicated in EMT [21, 22]. NF-ĸB acts as a significant player leading to the transactivation of EMT genes such as Twist, Snail, zinc finger E-Box binding homeobox 1 (ZEB1), ZEB2, and some EMT genes such as matrix metalloproteinase (MMP)-2 and MMP-9 [23–28]. There is ample evidence available suggestive of activation leading to the expression of mesenchymal genes by NF-ĸB in glioblastoma.

Emodin is a naturally occurring anthraquinone derivative containing hydroxyl groups at positions 1, 3, 8, and a methyl group at position 6. Emodin has been identified in 17 plant families globally so far [29]. Emodin has shown anticancer properties by inhibiting the phosphorylation of ERK 1/2, Akt/PKB, and FAK. In addition, it suppressed the transcription activity of two important transcription factors which include NF-ĸB and activator protein-1 (AP-1). Emodin induces apoptosis by G2/M phase arrest, decreased cell viability, increased ROS, Ca^2+^ production, and mitochondrial membrane potential loss [30]. It has also shown antioxidant, anti-inflammatory, anti bacterial and anti viral properties.

## 2. Materials and Methods

### 2.1. Protein Preparation

The coordinate file of PI3Kδ was procured from Protein Data Bank (PDB ID: 5UBT) [31]. Missing residues in the crystal structure were recovered using the SWISS-MODEL (https://swissmodel.expasy.org/interactive). Protein preparation, an important step to ensure structural correctness was performed using the Protein Preparation Wizard of Schrödinger package [32, 33]. Missing hydrogen atoms were added to the crystal structure and correct bond orders were assigned. This was followed by the filling of missing side chains and loops taking the advantage of Prime tool [34, 35]. Crystal waters were removed as per the standard protocol. Further, the redundant heteroatoms were deleted while keeping the co-crystallized ligand intact for subsequent use in grid specification [32, 36]. The structure was then optimized and water molecules having less than 3 hydrogen bonding interactions with non-water molecules were removed. Following this, minimization and subsequent grid generation were performed. To direct the incoming ligands toward the active site during the downstream molecular docking step the co-crystallized ligand was used as a centroid for grid generation [32, 36]. The grid-specified protein was then used as a receptor in molecular docking [34, 36].

### 2.2. Ligand Preparation

The 2D coordinates of 53 plant-derived small molecules (collected from literature) and the positive control Idelalisib were obtained from PubChem (https://pubchem.ncbi.nlm.nih.gov/). All 54 ligands were prepared using the LigPrep tool of the Schrodinger package [33, 36]. LigPrep enhances docking accuracy by removing errors in ligand molecules. Ligands were minimized, desalted and no metal binding states were generated. A computer program namely Epik was used for predicting the protonation states and the energy penalties of ligands [34, 37]. Thus, in nutshell, all the ligands were prepared under an identical set of parameters to restrain even minuscule fluctuations in docking score due to differential ligand preparation [32].

### 2.3. Molecular docking

After validating the docking algorithm using the self-docking protocol, we performed actual molecular docking using the Glide program (Glide v7.3). Prepared ligands (54) were docked against the grid-specified PI3Kδ receptor using the extra-precision flexible docking method [36, 38, 39]. The ligands were ranked based on docking score as Epik penalties are included in this score [34]. The docking scores were obtained from output pose viewer files and the most accurate pose of each ligand was taken into consideration [36].

### 2.4. Binding Free Energy Calculation

The binding free energy of docked complexes was calculated by using the Prime/molecular mechanics generalized born-surface area (MM-GBSA). This method employs an implicit solvation model for calculating free energies of solute-solvent interactions and molecular mechanics for measuring enthalpic contributions of protein-ligand interactions. Calculations were done in frozen conditions (receptor given no flexibility), using the VSGB solvation model and OPLS3 force field as previous studies have reported the best correlation with *in vitro* IC50 values under the defined receptor state [40–42]. This method performs several energy calculations from which binding free energy was calculated by using the below-mentioned equation;

ΔG (Bind) = Complex - (Receptor + Ligand)

More negative the value of binding free energy strong is the affinity of the ligand towards the receptor [32, 42].

### 2.5. Ligand interaction profile generation

The ligand interaction diagrams of the top 10 hits based on the evidence of docking score and binding free energy values were generated. The protein and a single ligand from the docked complex were imported into the workspace and the interaction profile was generated using default parameters [32]. Usually, more the number of interactions with ligands more favorable is the binding capacity of the inhibitor and vice versa [36, 43]. For the final selection of the top 3 hits, all three parameters including docking score, binding free energy value, and interactions towards PI3K delta receptor were taken into consideration.

### 2.6. 6 MD Simulation

GROMACS (Groningen Machine for Chemical Simulation) was used for the MD simulation, with the GROMOS96 43A1 force field. PRODRG web server was used to generate the calculated charges and topology files for the ligand. First, the system was relaxed enough to correct undesirable atomic contacts that may lead to unstable MD simulation. The system was hydrated by utilizing the Simple Point Charge 216 (SPC216) water model in a cubic simulation box. Appropriate counter ions either Na+ or Cl- were added randomly to achieve the net neutrality of the system. A free MD simulation of 10 ps with a force constant of 1000 KJ/mol/nm was performed during the equilibrium period. Subsequently, an equilibration was carried out at 300 K with isothermal-isochoric (NVT) using the modified Berendsen thermostat (v-rescale). Then the proposed isothermal-isobaric (NPT) algorithm was used to maintain the pressure of 1 bar. All bonds in the protein involving hydrogen atoms were constrained by the use of the LINCS algorithm. To illuminate the electrostatic interactions, the Particle Mesh Ewald (PME) method was used. To preserve the geometry of the water molecules SETTLE algorithm was used. Finally, the equilibrated system was taken for the final MD simulation for 100 ns simulation. Analysis including, root mean square deviation (RMSD), root mean square fluctuation (RMSF), and hydrogen bond analysis between protein and ligands was calculated with the help of GROMACS scripts. For graphical representation of the figures, OriginPro was used.

### 2.7. Reagents

Ham’s F12 media with 2 mM L-glutamine, 1.5 gms per liter sodium bicarbonate and Minimum Essential Medium Eagle (EMEM) with Earle’s salts, 2mM L-Glutamine, 1 mM Sodium Pyruvate, NEAA, 1.5 gms per liter sodium bicarbonate were obtained from HiMedia Laboratories Pvt ltd. Ttrypsin-EDTA solution, fetal bovine serum[FBS], penicillin-streptomycin, 3-(4,5-dimethylthiazol-2-yl)-2,5-diphenyltetrazolium bromide [MTT], dimethyl sulphoxide [DMSO], crystal violet, phosphoric acid sulfanilamide and Emodin (≥ 90 HPLC purified) were obtained from Sigma-Aldrich, Co. (St Louis, MO, USA). Lactate dehydrogenase [LDH] cytotoxicity assay kit was purchased from GBiosciences. Sybrgreen® mix was brought from Thermo scientific, the protein quantification assay kit from Bio-Rad, PVDF membrane was brought from Millipore-USA. All other chemicals were of ultrapure grade and obtained from Sigma-Aldrich, Co. (St Louis, MO, USA) or otherwise indicated.

### 2.8. Cell culture

The studies were performed on the C6 cell line (Rattus Norvegicus glioma-derived cell line) and U87-MG (Homo sapiens-derived cell line). The cells were cultured in Ham’s F12 media and EMEM media respectively supplemented with 10% FBS and 100 units of penicillin-streptomycin and maintained at 37°C in a humidified incubator with a 5% CO2 atmosphere. All the experiments reported here were performed under the protocol as: cells were grown to 75-85% confluency, trypsinized, and subcultured in culture dishes at a concentration of 2×10^4^ cells/ ml. The cells were continually maintained in a humidified incubator with 5% CO2 at 37°C.

### 2.9. Preparation and recovery of liquid nitrogen cell stocks

Liquid nitrogen-stored cell stocks were prepared by trypsinizing 70-80% confluent cell cultures and centrifuging the dissociated cells (1000-2000 rpm, 4-5 min at room temperature). Cell pellets were resuspended in a medium containing 90% FBS and 10% DMSO. 1 ml aliquots were pipetted into cryotubes, which were brought to −80°С slowly. The cryotubes were then placed in liquid nitrogen for long-term storage. Cell stocks were recovered by thawing for 1 min in a water bath at 37°С. These cells were seeded in a fresh sterile T-25 flask together with pre-warmed culture medium and grown under standard growth conditions.

### 2.10. MTT cell proliferation assay

Cell proliferation measurement forms the basis for evaluating the effects of drugs on cells *in-vitro.* Mitochondrial dehydrogenase enzymes of metabolically active cells reduce MTT-tetrazolium to generate reducing equivalents and aqueous insoluble intracellular purple formazan crystals. Aqueous insoluble formazan crystals formed are dissolved in DMSO or an acidified solvent, which is then quantified spectrophotometrically. The effect of emodin and Rutin on the proliferation of C6 and U87 cells over a range of concentrations were quantified using MTT colorimetric assay. For each condition, 2×10^4^ cells/well were seeded in 24 well plates in triplicates and allowed to grow. Media was changed after 24 h and cells were treated with drugs in increasing concentration. The plates with media and controls were incubated for 24 hr at 37°C and 5% CO2. MTT was prepared by dissolving in PBS at 5 mg/ml. After 24 h, culture media with drugs was carefully removed and 500μl MTT solution was added and plates were incubated for 2 h at 37°C. The supernatant containing MTT solution was discarded and MTT solvent (4 mM HCl and 0.1% NP-40 in isopropanol) was added to all wells. The plates were placed on an orbital shaker at room temperature for 20 min in dark to mix the contents thoroughly and dissolve the blue formazan crystals trapped inside the cells. After dissolution, the plates were read on an ELISA plate reader (BioTek instruments inc.) at a reference wavelength of 650 nm and test wavelength of 560 nm.

### 2.11. Lactate dehydrogenase cell death assay

LDH assay was performed by evaluating soluble cytosolic LDH enzyme release in the culture media as a cell death marker. We utilized the CytoScanTM LDH Cytotoxicity Assay kit (G Biosciences) to assess the effect of the drug on cell viability. LDH is a mitochondrial enzyme that gets released from dead cells into the cytosol where it reduces NAD+ to NADH in the presence of L-lactate. The NADH formed can be measured in a coupled reaction wherein tetrazolium salt is reduced to a red formazan product. The soluble and highly colored formazan produced is quantitated spectrophotometrically at 490nm. In brief, 2 × 10^4^ C6 cells/ml were grown in 24 well plates and in addition to test samples, negative and positive controls were also included. The media was changed after 24 h and cells were treated with an IC-50 dose of emodin. During the experiment, 100 μl of culture media was aspirated at four different time points (6 h, 12h, 24h, and 48 hr) and transferred to 96 well plates. To each well containing 100 ul of media, 100 μl of LDH substrate was added and the mixture was incubated for 20 min at RT. The reaction was stopped by adding 50 μl of CytoScanTM stop solution. LDH activity of samples was measured spectrophotometrically at 490 nm by ELISA plate reader (BioTek instruments inc.). LDH concentrations were calculated by setting standards along with experimental samples.

### 2.12. Quantification of apoptosis by propidium iodide staining

Propidium iodide (PI) passes through the membrane of damaged/ dead cells and intercalates into dsDNA in a stoichiometric manner, thus acts an apoptotic indicator stain. The permeability of the membrane is altered when a cell suffers damage and its membrane integrity is lost. PI moves into the cell and stains DNA thereby signaling the extent of membrane damage and thus cytotoxicity. C6 were seeded at a density of 2 × 10^4^ cells/ml in complete medium in six-well plates. The cells were treated with emodin for 12 h and 24 hr along with respective controls. After respective periods, the medium was aspirated from corresponding wells, and cells were washed gently twice for 5 min each time with prechilled PBS. Washing was followed by incubating cells with PI solution (5 μg/ml) for 30 min in dark. Then the cells were washed with TBS (50 mM Tris-HCl, pH 7.3). For determining the level of background signal, wells without cells having PI only were taken as control. The background signal of dead cells was determined from cells treated with 0.2% Triton X-100 for 10-15 min. To avoid chances of changes in dead and living cell distribution as a result of the staining/ washing procedure, the temperature of samples was maintained at 4°C. Wells containing stained cells were visualized at Floid Cell Imaging Station (Life Technologies) and by measuring fluorescence signal at excitation/emission wavelength of 493/636 nm using a spectrofluorophotometer (Shimadzu, RF-5301). Six random fields were chosen for observation.

### 2.13. Apoptotic analysis by Annexin V (FITC) Flow Cytometry

Cells undergoing apoptosis expose phosphatidylserine on the outer leaflet of cells which has a high affinity for fluorescently labeled Annexin V. There occurs a difference in fluorescence intensity between non-apoptotic and apoptotic cells stained with annexin V conjugates which are measured by flow cytometry. Cells were grown and treated for 6 h, 12 h, and 24 h in 12 well culture plates. Next, adherent and floating cells were harvested and washed with ice-cold PBS twice. The washed cell samples were resuspended in 500ml of binding buffer containing 3 ml Annexin-V for 10 mins and 2 ml 7-AAD for 15 mins in dark. These samples were subsequently evaluated for apoptosis. Flow cytometry was performed at the Department of Biotechnology, National Institute of Technology, Rourkela Odhisa, India on a BD Accuri TM C6 Flow Cytometer. Apoptotic events were expressed as the percent of sub-G1 cells or the percentage of apoptotic cells (Combining early apoptotic Annexin V+ /7-AAD- and late apoptotic Annexin V+/7-AAD+ cells).

### 2.14. Reactive Oxygen Species analysis

ROS production at a low level is a fundamental feature of living cells. At low levels, it plays an important role in various signaling transduction pathways. However, higher levels damage proteins, nucleic acids, and lipids under oxidative stress. Elevated levels of ROS during oxidative stress are associated with apoptosis/ necrosis, aging, and various other pathological conditions such as diabetes, vascular disease, pulmonary disorders, and cancer. ROS measurements were performed using DCFDA/H2DCFDA kit (Abcam). C6 cells were harvested, seeded at a density of 2.5 x10^4^ cells per ml in complete medium in six-well plates, and allowed to attach overnight. After overnight incubation, the medium was aspirated and cells were washed once in 1x buffer. Cells were stained with 25μM DCFDA in 1x buffer for 45 mins at 37°C. After incubation, wash cells once in 1xPBS/1x Buffer. Add 100 μL/well of treatment compound and incubate for 2 h, 4 h, and 6h time points. After this, the fluorescence signal was measured at an excitation wavelength of 485nm and an emission wavelength of 535nm using a spectrofluorometer (Shimadzu, RF-5301). Changes in the percentage of control were measured after background subtraction.

### 2.15. Protein extraction

Cells were seeded in 60-mm culture dishes and allowed to grow for 24 h. After this, the culture medium was removed and cells were treated with the drug for appropriate periods. Next, the culture medium was removed and cells were washed twice with pre-chilled PBS and scraped into NP-40 lysis buffer (137 mM NaCl, 20 mM Tris-HCl (pH 8), 1% Nonidet P-40, 10% Glycerol, 1mM PMSF, 5-10 mM NaF, 2 mM EDTA, and phosphatase inhibitors). Scraped cells were incubated on ice for 45-60 min and centrifuged at 12,000 rpm for 12 mins. The supernatant was poured into fresh pre-chilled Eppendorf and samples were stored at −80°C till further use.

### 2.16. Protein estimation

Protein estimation was done using Bradford assay

### 2.17. Western blotting

Proteins separated on the gel were transferred to PVDF membrane by semi-dry transfer process following manufacturer’s instructions (Hoefer-USA) for 1-2 hours at 200-400 mA. After transfer, the PVDF membrane was blocked with 5% BSA in PBS for 1 h at room temperature. Then, the membrane was washed once with PBST (PBS containing 0.5% Tween-20) and incubated overnight with specific primary antibodies: [AKT (Pan) (1:2000), Phospho-Akt (Thr308)(1:1000), IĸBα (1:1000), NF-ĸB p65 (1:1000), N-Cadherin (1:1000), β-Catenin (1:1000), Claudin-1 (1:1000), Histone H3 antibody (1:1000) cell signaling technologies, Danvers, Massachusetts, USA] β-actin (1:200-500) (Santa Cruz Biotechnologies Dallas Texas, US). After incubating the membrane overnight with the corresponding antibody at 4°C, the membrane was washed three times for 10 mins each time with PBST. The detection was performed by secondary antibody using anti-rabbit IgG DyLight® 800 conjugate (1:10,000) or mouse IgG DyLight® 680 conjugate (1:10,000) and the membrane was incubated in corresponding secondary antibody for 2 hours at room temperature. After incubation, the membrane was washed thrice for 10 mins with PBST and was then detected using Odyssey infrared detection system (LI-COR). Band intensity was calculated from scanned blots using ImageJ software. β-actin and H3 antibodies were taken as loading control for cytoplasmic and nuclear proteins respectively and normalization with corresponding control antibodies.

### 2.18. Preparation of nuclear and cytoplasmic fractions

The cells were washed two times with PBS. Afterward, cells were scraped in fresh PBS, and contents were poured into 15 ml falcon. The cells were centrifuged at 3000 rpm for 5 mins and the supernatant was discarded. The pellet was then resuspended in 100 μl/10^6^ cells of 1x Pre-extraction buffer and transferred to a 1.5 ml micro-centrifuge tube and incubated for 10 mins on ice. It was followed by vigorous vortexing of the tube for 10 seconds. Then, the tube was centrifuged for 1 min at 12,000 rpm. After this step, the separated cytoplasmic fraction can be carefully removed from the nuclear pellet. For extraction of nuclear protein, DTT solution and PIC were added to the extraction buffer in a 1:1000 ratio. Two volumes of Extraction Buffer containing PIC and DTT were added to the nuclear pellet (∼ 10 ul / 10^6^ cells). The preparation was then incubated on ice for 15 mins with intermittent vortexing (5 seconds after every three mins). The suspension was centrifuged at 14,000 rpm for 10 mins at 4°C and the supernatant was then transferred to a fresh microcentrifuge tube. The protein concentration of both fractions was quantified using the Bradford method.

### 2.19. Human Phospho-kinase Array kit

Cells were seeded in 60 mm dishes and allowed to grow till 70-80% confluency is attained which was followed by medium change and drug treatment for 30, 45 and 60 mins. After a specific time, lysates of specific conditions were collected and the relative phosphorylation status of 37 kinase phosphorylation sites was evaluated by using a Human Phospho-kinase Array kit (ARY003C, R&D Systems, Minneapolis, MN, USA) following the manufacturer’s instructions. Briefly, the kit is a membrane-based sandwich immunoassay. On the nitrocellulose membrane provided, antibodies are captured and spotted in duplicates which bind to corresponding proteins there in the sample. The membranes are incubated with cell lysates overnight on a rocking platform at 4°C. The membranes were then washed three times for 10 mins each time to remove unbound proteins. Next, phosphorylation levels of bound proteins were detected by biotinylated phospho-specific antibodies. The detection cocktail comprising of streptavidin–HRP and chemiluminescent reagent were applied adequately to produce a signal at captured spots which correspond to the amount of bound phosphorylated protein. The chemiluminescent signal produced was acquired by the documentation system Epson Expression 11000xL and was analyzed using ImageJ software. The pixel density of each spot was calculated following subtraction of the background (negative control) and normalization to the positive control spots.

### 2.20. RNA isolation and quantitative Real-Time-PCR assay

Total RNA was isolated from C6 cells using TRIzol® reagent (*Invitro*gen, USA) according to the manufacturer’s instructions. The integrity and quantity of RNA were determined by electrophoresis of total RNA through 1 % denaturing agarose gel. Expression of EMT markers such as N-Cadherin, β-Catenin, Claudin-1, Slug, Snail, Vimentin, and Zeb1 was evaluated by qRT-PCR assay which is based on SYBR® Green gene expression analysis detection system (Thermo Scientific). For loading control, we used β-actin. We designed gene-specific forward and reverse primers through NCBI Primer blast software (Table 1). After carrying out DNase treatment on isolated RNA, we performed reverse transcription using a standard protocol. cDNA first strand was generated by using MMLV reverse transcriptase (Thermo Scientific) and random hexamers in a reaction mixture of 20 μl containing 5μg of total RNA. PCR amplification was carried out in Benchmark DNA thermal cycler (TC 9639). The reaction parameters for the synthesis of cDNA were as follows: 25.0°C, 7 min; 40.0°C, 60 min; 65°C, 15 min. The quality of cDNA and primer specificity to be used in qRT-PCR was evaluated using end-point PCR with Taq DNA polymerase. Expression levels of all genes concerned were determined using specific primer pairs for the genes with β-actin as a loading control in qRT-PCR analysis. We amplified 3.5 μl of synthesized cDNA in Real-Time PCR 7500 (Applied Systems) using maxima Sybrgreen® mix according to the manufacturer’s instructions. qRT-PCRs were performed in a MicroAmp® optical 96-well plate (Applied Biosystems). All the reactions were run in triplicates as follows: 94°C, 30 s; 60°C, 30 s; 72°C, 90 s; 40 cycles, finally extending at 72°C for 5 min.

**Table 1:**
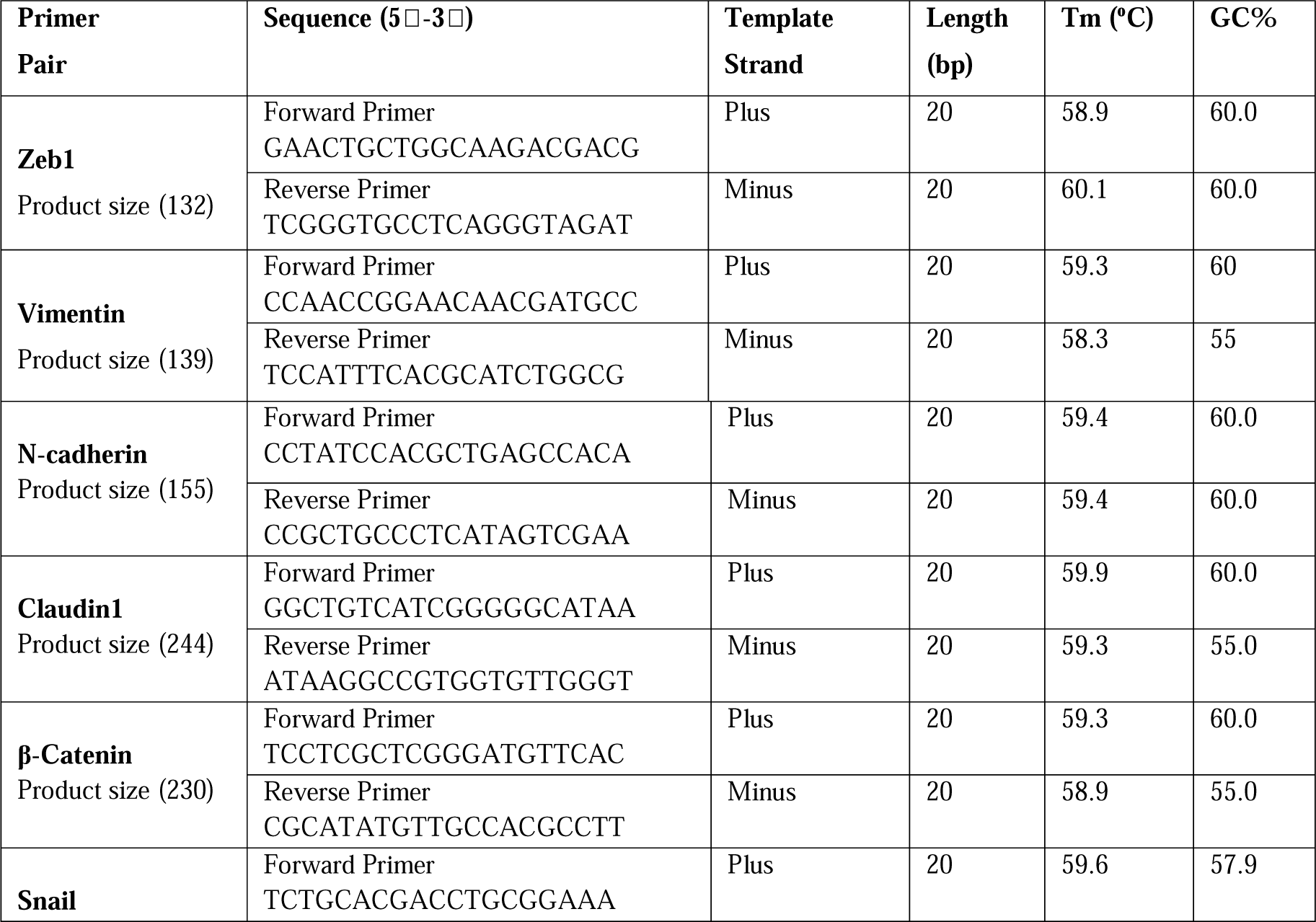
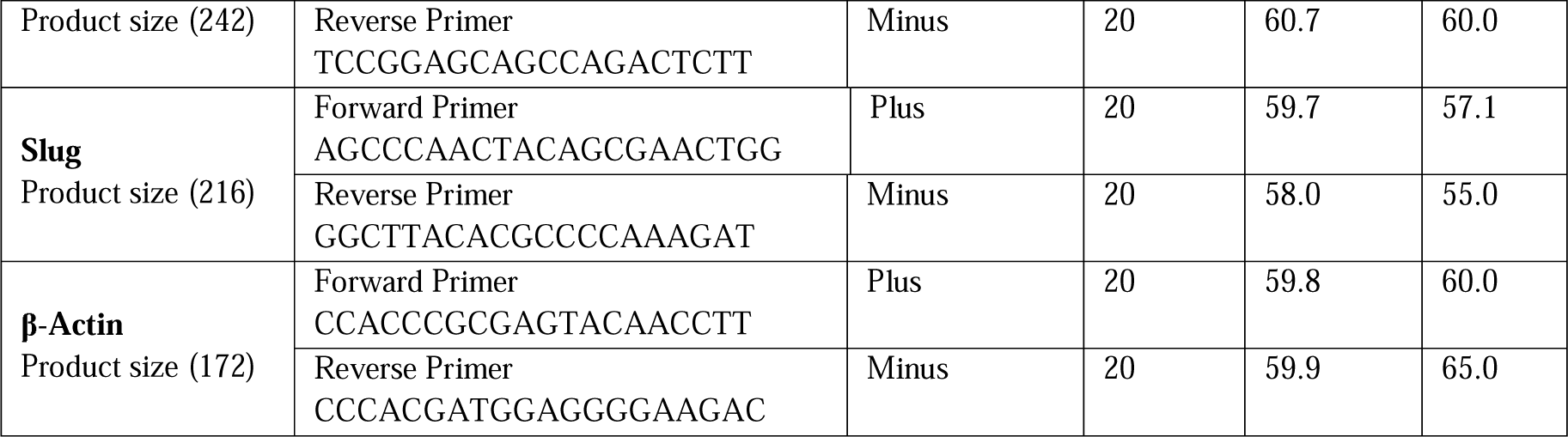
Sequence and characteristic properties of gene specific forward and reverse primers for respective gene amplification.

### 2.21. Confirmation of product and primer specificity

Cycle threshold [Ct] value, the point at which PCR product is first detected above the threshold, was evaluated for each sample and the average Ct of triplicate samples was determined. Melt curve analysis was used in the range of 60–95°C using Real-time PCR. Besides, PCR product length was confirmed by electrophoresis on 2% agarose gel.

### 2.22. Quantification of expression of target genes

To quantify target gene-specific transcripts in drug-treated cells, the corresponding Ct values were normalized first by subtracting the Ct value acquired from β-actin control. The concentration of gene-specific mRNA in treated cells relative to untreated cells was calculated by subtracting the normalized Ct values obtained for untreated cells from those obtained for treated cells and the relative concentration was determined.

## 3.0 Results

### 3.1. In-Silico Analysis

We adopted an in-silico approach to screening a range of natural molecules for a potent P110δ inhibitor based on the structural similarity of Idelalisib/Cal-101, which is a synthetic P110δ inhibitor approved for use. We carried out protein preparation, ligand preparation, molecular docking, calculation of binding free energy, and ligand interaction profile generation.

For protein preparation, a coordinate file of PI3Kδ was procured from the protein data bank, and missing residues in the crystal structure were recovered. Missing hydrogen atoms, missing side chains, and loops were added and the crystal water was removed from the structure appropriately. Following this, minimization and subsequent grid generation were performed. To direct the incoming ligands toward the active site during the downstream molecular docking step, the co-crystallized ligand was used as a centroid for grid generation. The grid-specified protein was then used as a receptor in molecular docking. For ligand preparation, 2D coordinates of 53 plant-derived small and the positive control Idelalisib were obtained from PubChem. Ligands were minimized, desalted and no metal binding states were generated. All ligands were prepared using an identical set of parameters to restrain even small fluctuations in docking score due to differential ligand preparation. Following protein preparation, prepared ligands were docked against the grid-specified PI3Kδ receptor and ranked according to the docking score. It was followed by calculating binding free energies, for “more negative the value of binding free energy, strong is the affinity of the ligand towards the receptor.” The ligand interaction diagrams of the top 10 hits based on the evidence of docking score and binding free energy values were generated. The protein and a single ligand from the docked complex were imported into the workspace and the interaction profile was generated using default parameters. Usually, the number of interactions with ligands more favorable is the binding capacity of the inhibitor and vice versa. For the final selection of the top 2 hits, all three parameters including docking score, binding free energy value, and interactions towards PI3K delta receptor were taken into consideration (Table 2). Figure 1, 2 and 3 represent ligand interaction diagrams of Idelalisib, Emodin and Rutin respectively.

**Figure 1:**
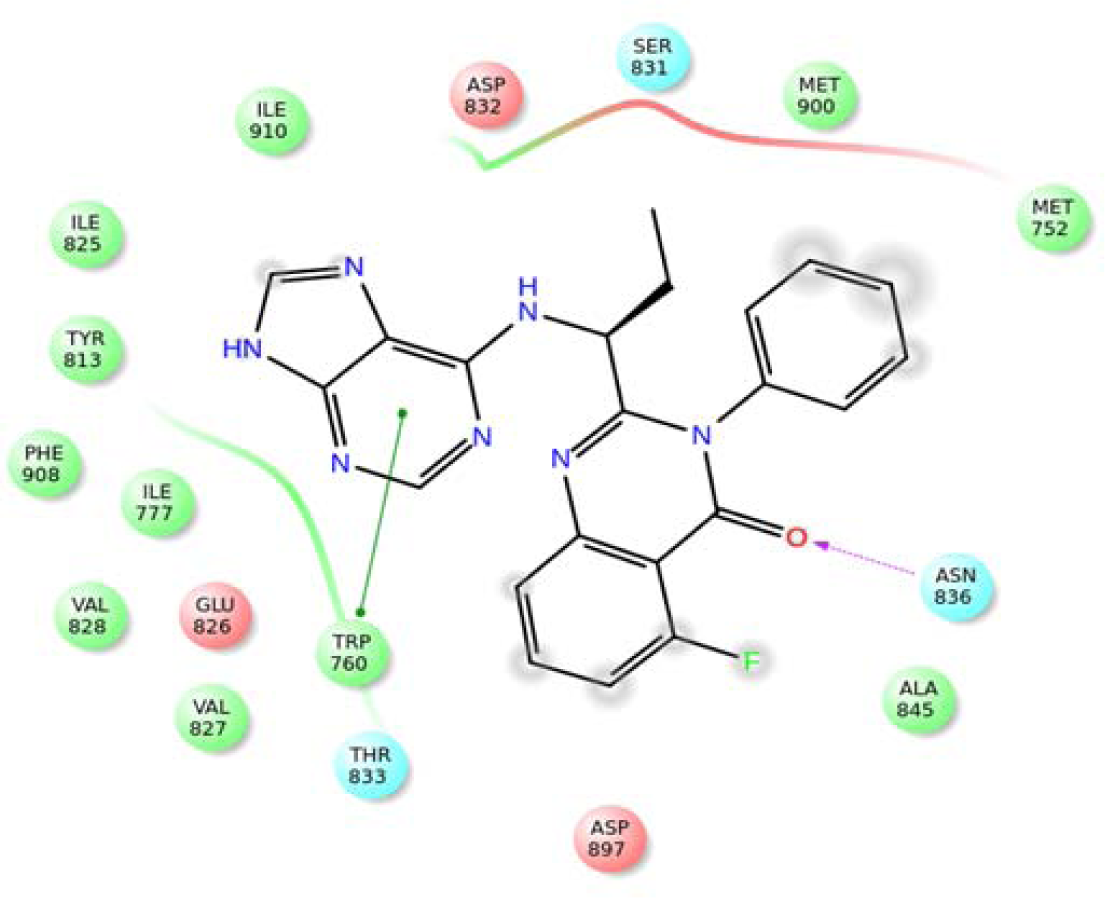
Ligand interaction diagram of idelalisib.

**Figure 2:**
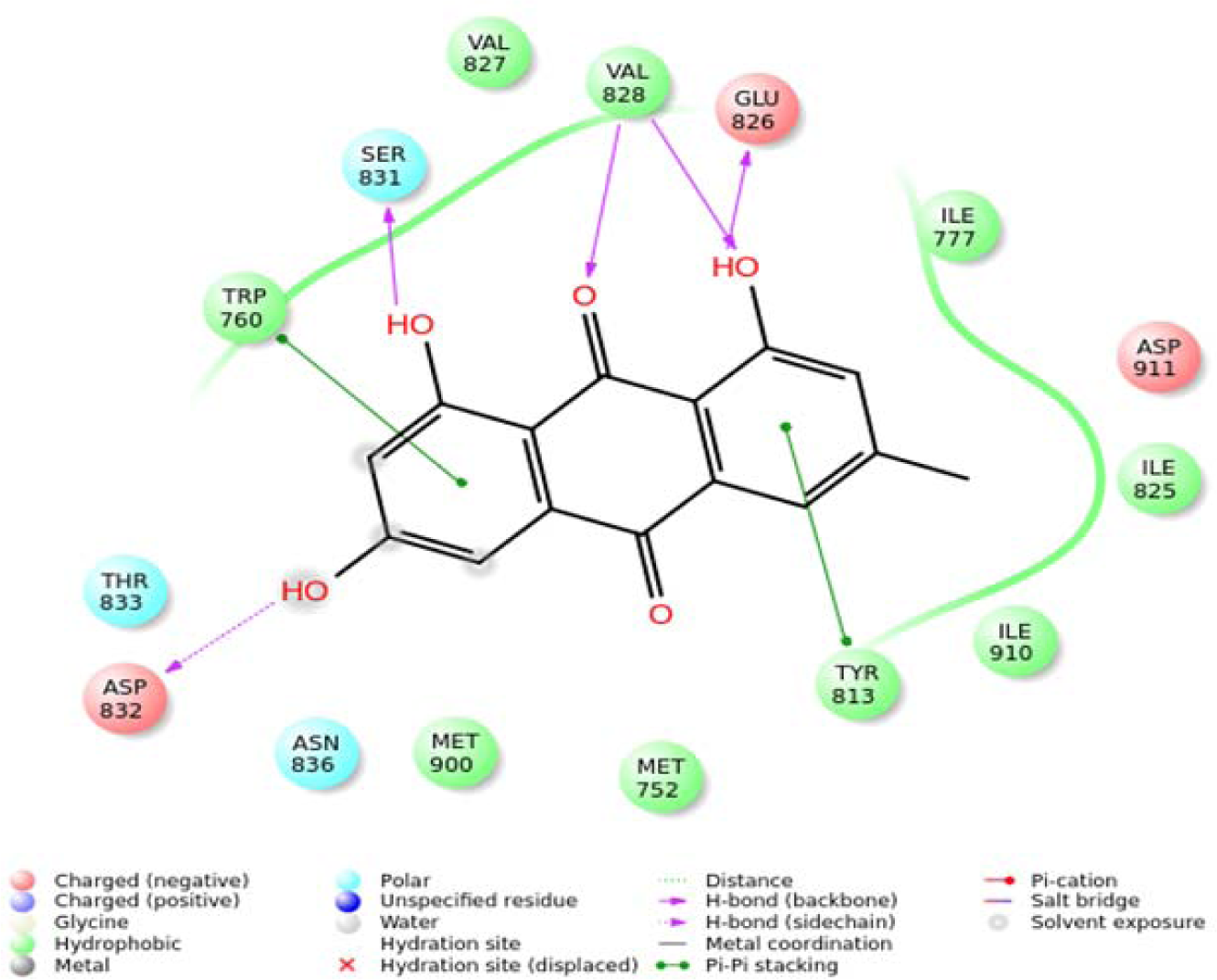
Ligand interaction diagram of Emodin.

**Figure 3:**
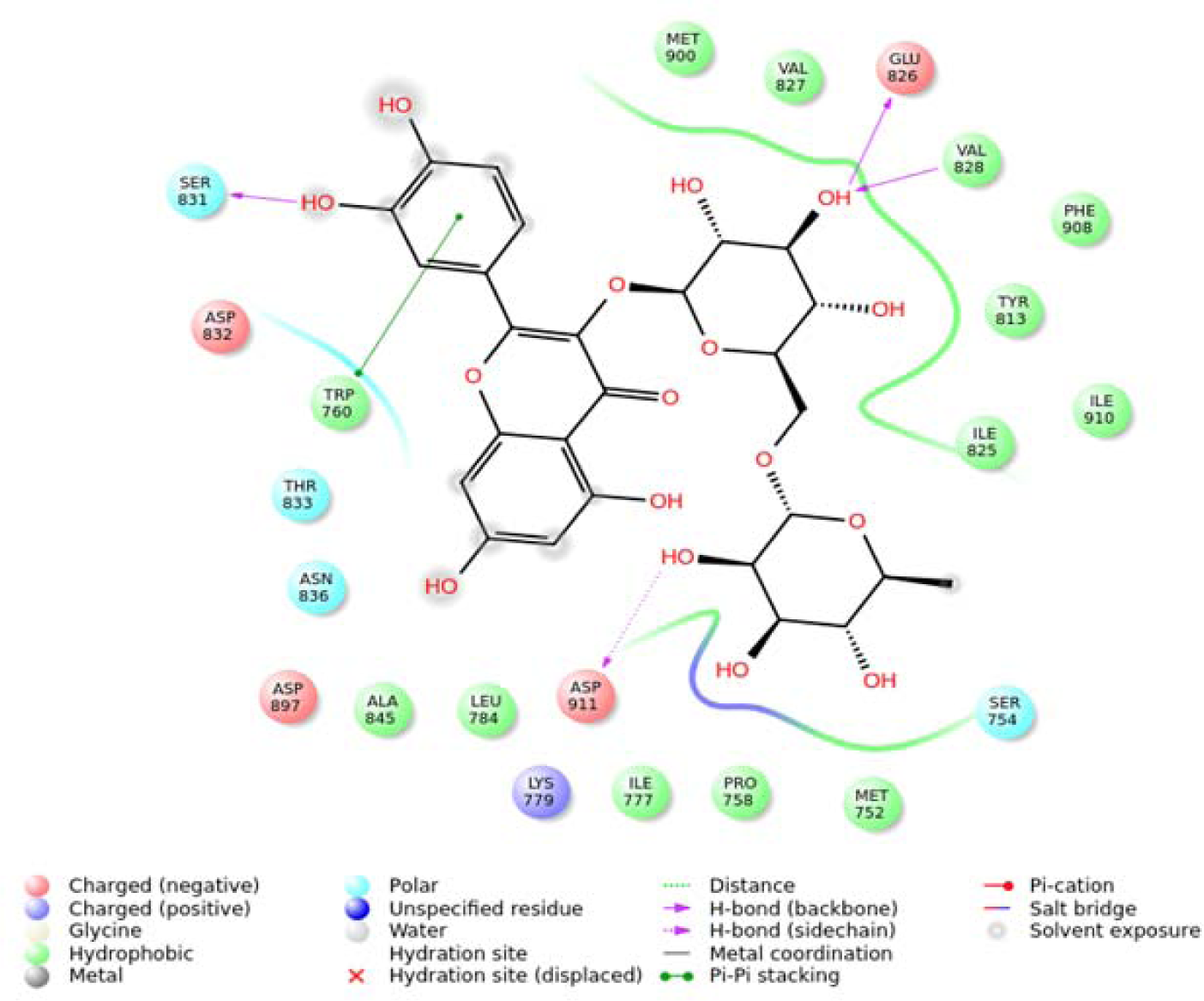
Ligand interaction diagram of rutin.

**Table 2:** Pubchem Id, Docking score and Delta G binding details of Idelalisib (Reference molecule), Rutin and Emodin.

| S.No | Common Name | Pubchem Id | Docking Score | Delta G binding |
| --- | --- | --- | --- | --- |
| 1 | Idelalisib | 11625818 | -6.226 | -26.9117 |
| 2 | Rutin | 5280805 | -11.821 | -53.9549 |
| 3 | Emodin | 3220 | -11.362 | -43.2643 |

### 3.2. In-vitro Analysis

C6 and U87-MG cells were cultured in Ham’s F12 and Eagle’s Minimum Essential media respectively. Both media were supplemented by 10% FBS and penicillin/streptomycin antibiotic. The cells were grown in a humidified incubator with having 5% CO2 level at a temperature of 37°C. Sub-culturing of cells was done as per experimental requirement in 6, 12, and 96-well plates. The cells were seeded at a density of 2×10^4^ cells/ ml and were grown to 70-80% confluency following experimental requirements. All the results are representative of three independent experiments.

### 3.3. Treatment of cells with Emodin significantly reduces the proliferation of glioma cells

The molecules shortlisted based on docking score and Delta G binding energies were assayed for their effect on cell proliferation and dose standardization. To evaluate the effect of emodin and rutin on cell proliferation in C6 and U87-MG cell lines MTT assay was done as described in the materials and methods section. U87-MG and C6 cells were treated with emodin over a concentration range of 30 μM to 300 μM and incubated for 24hr. The proliferation of both cell types decreased by treating them with increasing doses of emodin in comparison to untreated control cells. A 50% decrease in cell proliferation was observed at 120 μM in U87-MG as well as C6 cells (Fig 4 and Fig 5), for further studies with emodin, this concentration was used. Similarly, U87-MG and C6 cells were treated with Rutin for 24hr over a concentration range of 200 μM to 2mM. The proliferation of both cell types decreased by treating them with increasing doses of Rutin in comparison to untreated control cells. There was about a 50% decrease in cell proliferation at 1.2 mM concentration in both the cell types (Fig 6 and Fig 7), for further studies with rutin, we used this concentration of rutin.

**Figure 4:**
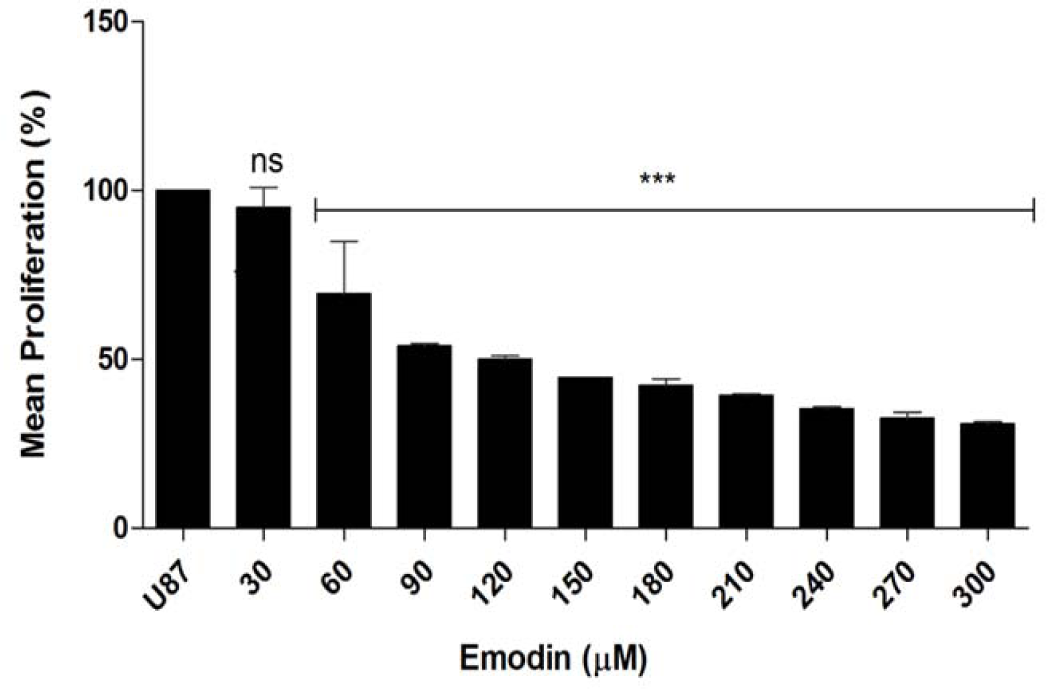
Dose responsiveness of U87 glioma cells treated with the indicated concentrations of Emodin (30μM t 300 μM) as assessed by MTT assay: Emodin inhibited proliferation of U87-Mg cells in a dose dependent manner, IC-50 being 120 μM. Data are expressed as mean ± SD. Bars indicate standard deviation and asterisk represent significance *P < 0.05; **P< 0.01; ***P< 0.001.Dunnett’s Multiple Comparison Test was used to analyze the data.

**Figure 5:**
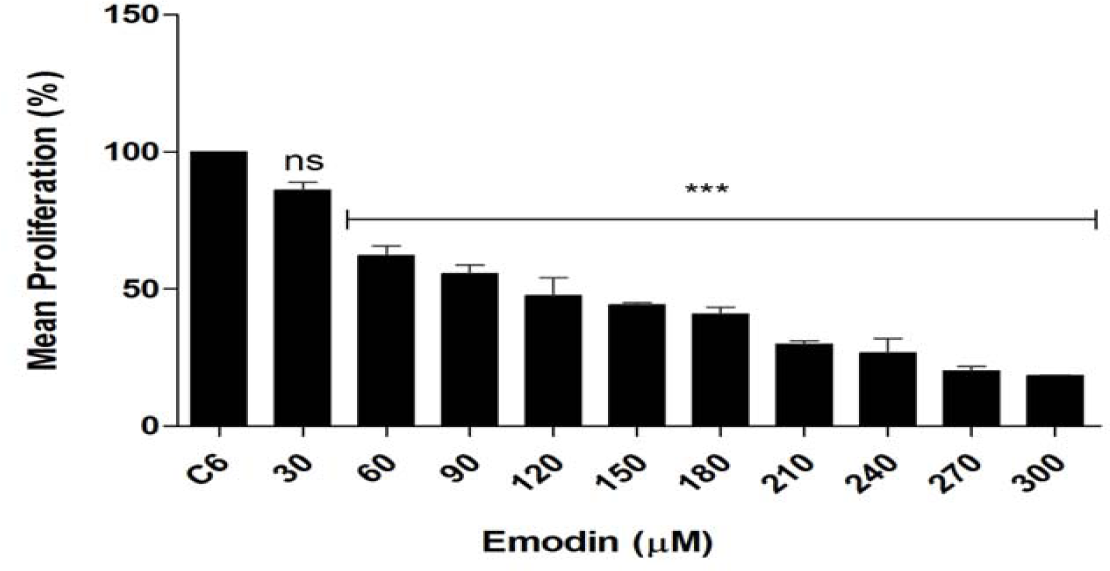
Dose responsiveness of C6 glioma cells treated with the indicated concentrations of Emodin (30 μM to 300 μM) as assessed by MTT assay: Emodin inhibited proliferation of C6cells in a dose dependent manner, IC-50 being 120 μM. Data are expressed as mean ± SD. Bars indicate standard deviation and asterisk represent significance *P < 0.05; **P< 0.01; ***P< 0.001.Dunnett’s Multiple Comparison Test was used to analyze the data.

**Figure 6:**
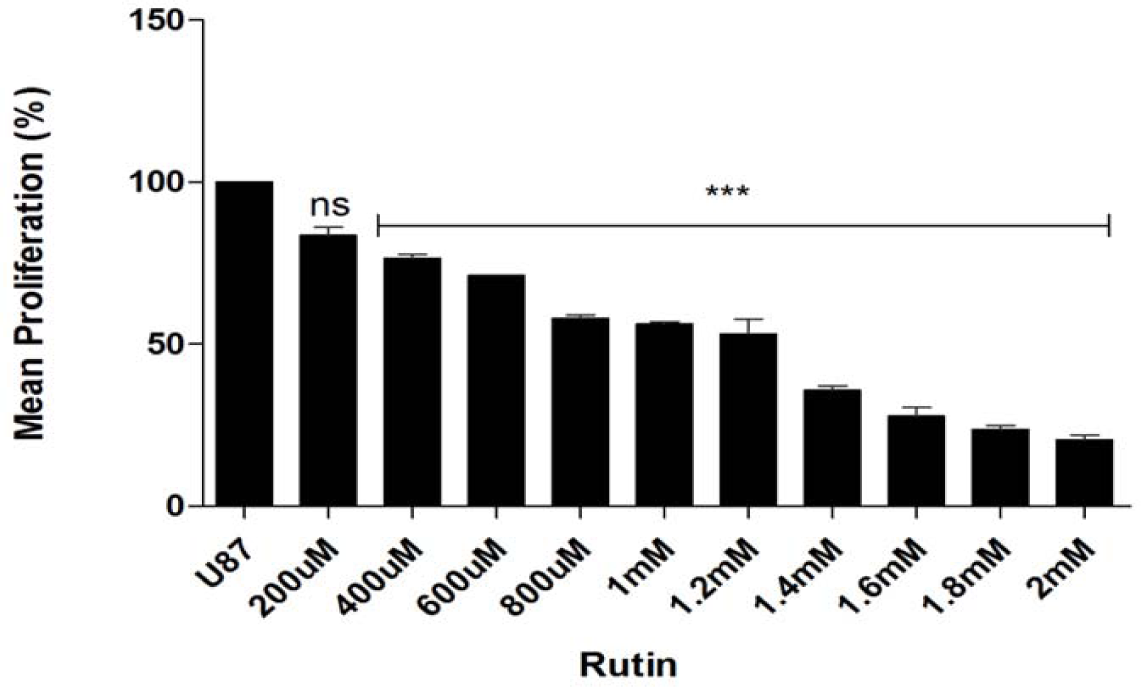
Dose responsiveness of U87-MG glioma cells treated with the indicated concentrations of Rutin (20 μM to 2mM) as assessed by MTT assay: Rutin inhibited proliferation of U87-MG cells in a dose dependent manner, IC-50 being 1.4mM. Data are expressed as mean ± SD. Bars indicate standard deviation and asterisk represent significance *P < 0.05; **P< 0.01; ***P< 0.001. Dunnett’s Multiple Comparison Test was used to analyze the data.

**Figure 7:**
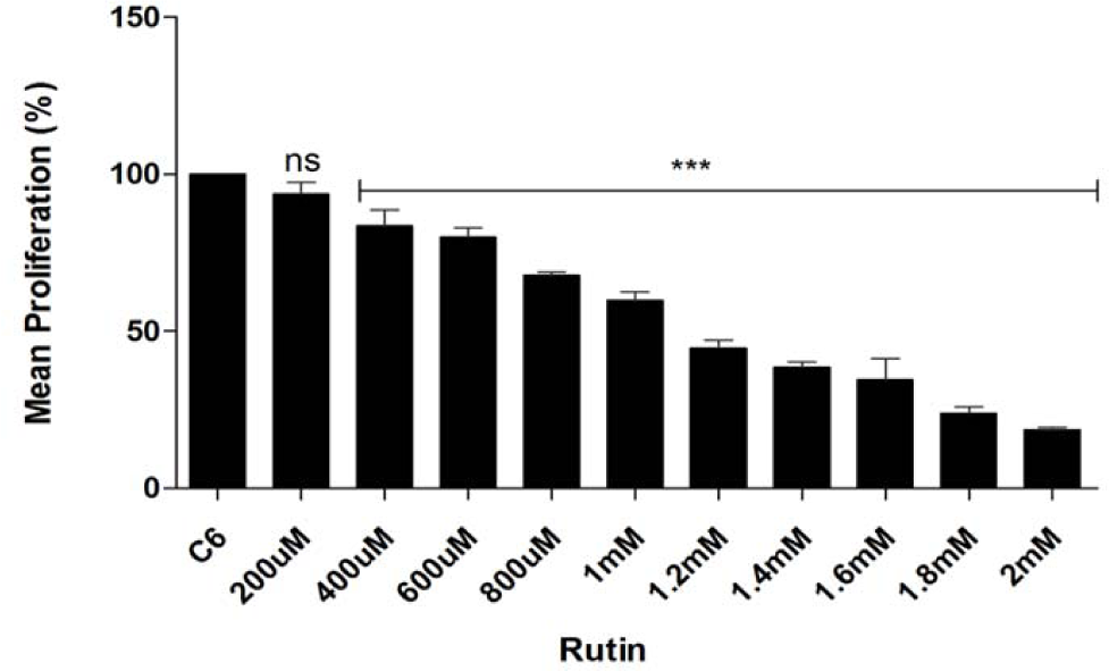
Dose responsiveness of C6 glioma cells treated with the indicated concentrations of Rutin (200 μM M to 2mM) as assessed by MTT assay: Rutin inhibited proliferation of C6 cells in a dose dependent manner, IC-5 being 1.4mM. Data are expressed as mean ± SD. Bars indicate standard deviation and asterisk represent significance *P < 0.05; **P< 0.01; ***P< 0.001. Dunnett’s Multiple Comparison Test was used to analyze the data.

### 3.4. Emodin enhances glioma cell death

We further examined the effect of emodin treatment on cell death in glioma cells. It was accomplished by LDH assay which depicts plasma membrane damage by cytotoxic agents. Treatment of C6 glioma cells with 120 μM concentration of emodin caused ∼35% cell death after 6hr, ∼55 % after 12hr, ∼85 % after 24hr and ∼95% cell death after 48hr compared to control (Fig 8).

**Figure 8:**
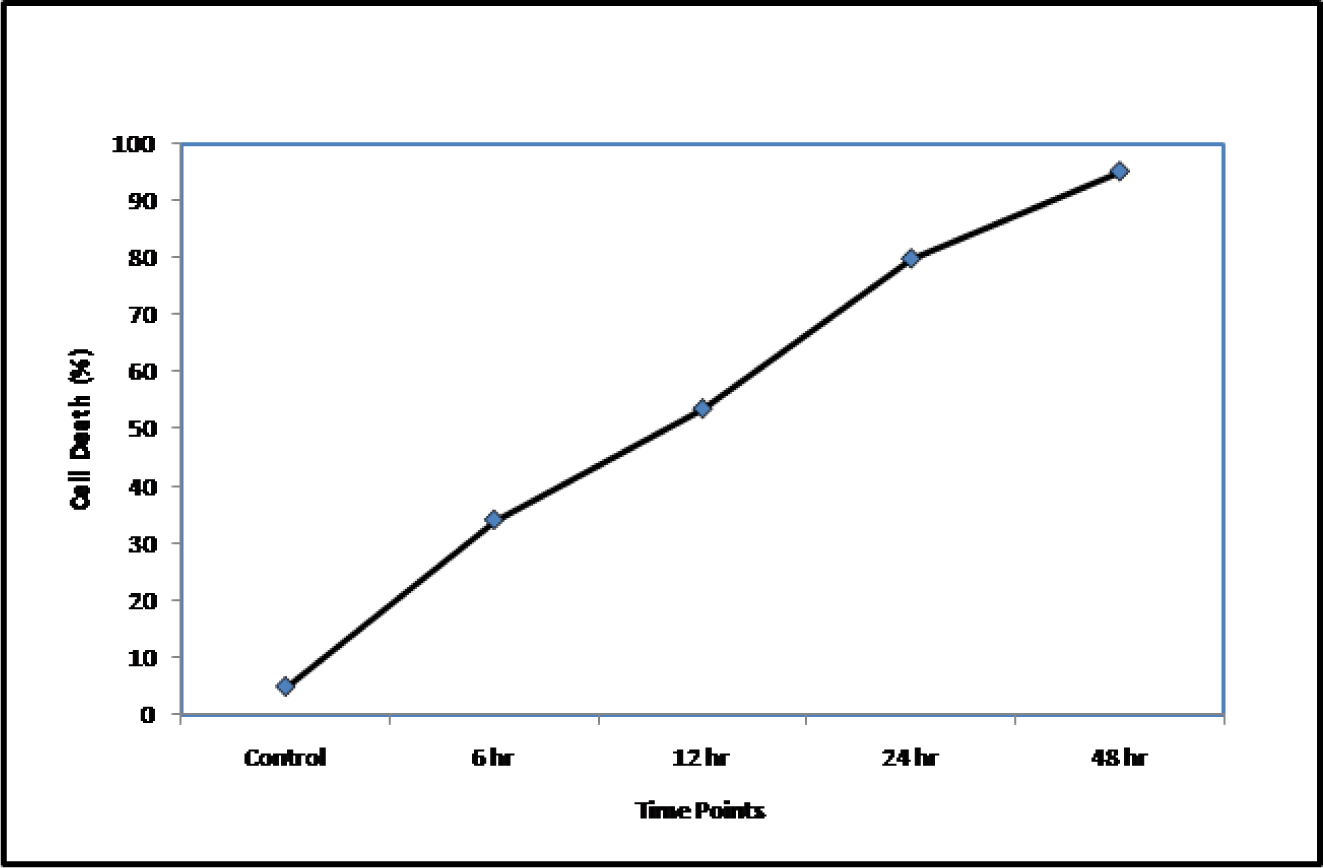
Effect of Emodin (120μM) on viability of C6 cells at selected doses and at different time point’s in comparison to control cells: Treating C6 glioma cells with 120 μM concentration of emodin caused ∼30% cell death after 6hr, ∼50 % after 12 hr, ∼80 % after 24hr and ∼ 95% cell death after 48hr compared to control cells.

### 3.5. Emodin induced cell damage accompanied by apoptosis

We performed PI staining which is an indicator of cell damage and apoptosis under non-fixed cell staining conditions thus differentiating stress induced morphological changes from apoptosis induced ones. Results showed that the treatment of C6 cells with emodin significantly induced cell apoptosis in comparison to control group. In control group after 12 hr and 24 hr, nuclei stained with PI accounted for about ∼ 4.5% and ∼ 17.5% respectively, while as ∼ 37.5% and ∼92% for emodin treated cells respectively (Fig 9A). Spectroflourometric analysis also show similar pattern in apoptotic results as depicted in Fig 9B. These findings reveal morphological changes characteristic of apoptosis.

**Figure 9:**
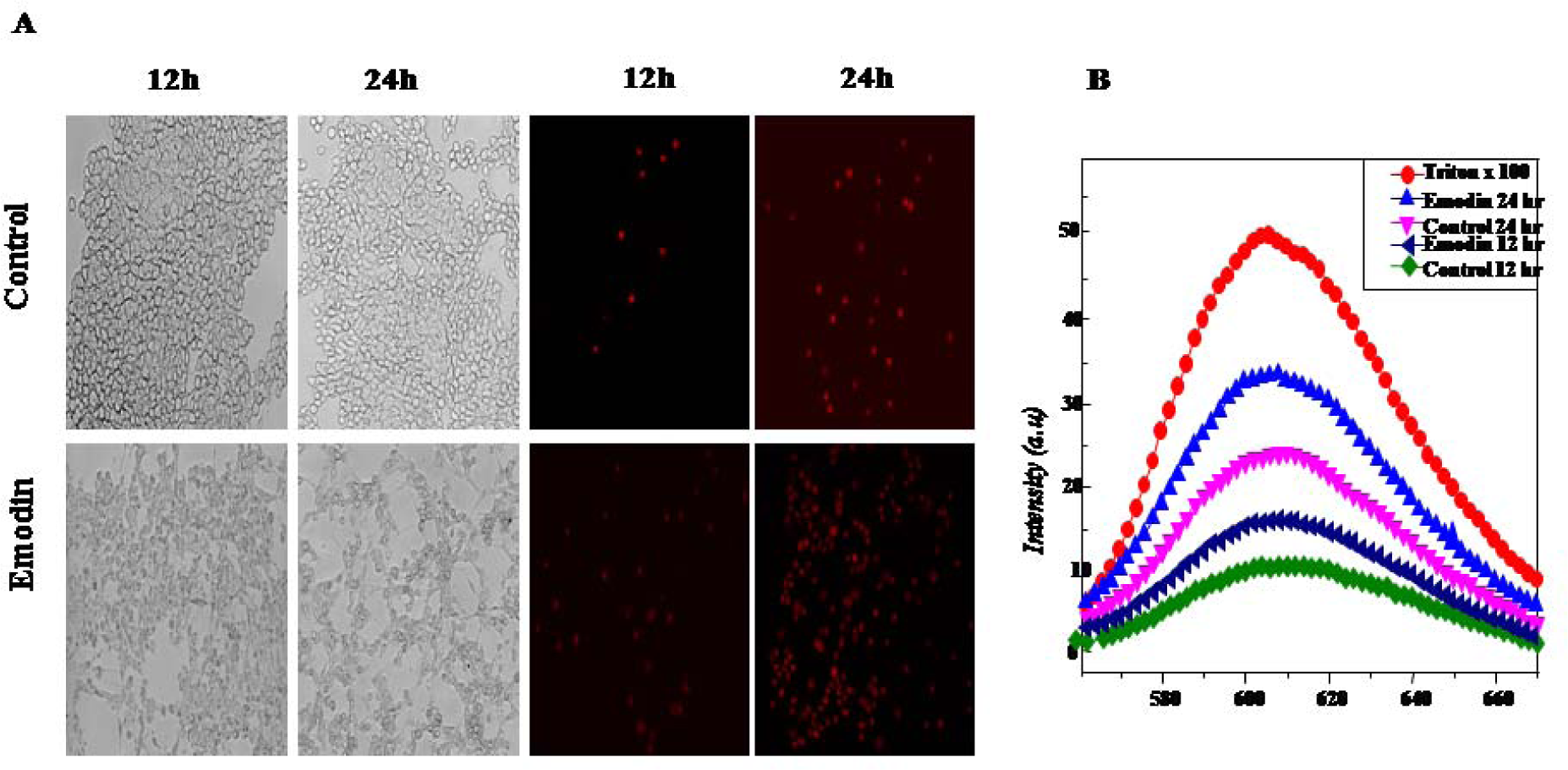
Photomicrographs depicting, black& white and PI stained fluorescent images (A; as an index of cell damage (scale bar: 100μm) (B)corresponding spectrofluorometer images: Emodin treated group showed larger number of nuclei stained with PI after 12hr and 24hr as compared to control.

### 3.6. Emodin induces apoptosis in C6 glioma cells

Apoptosis induction in C6 glioma cells was evaluated by Annexin V-FITC flow Cytometry. The 24 h incubation of cells with emodin resulted in a gradual increase in the fluorescence of FITC bound cell population which directly implied increase of apoptotic cells. In control group of cells, total apoptotic cell population accounted for ∼6.64%, ∼8.6% and ∼8.7% after 6 hr, 12 hr and 24 hr of incubation, however total apoptotic cell population after emodin treatment accounted for about ∼25.6%, ∼23% and ∼20.5% at indicated time points respectively (Figure 10A and 10B). This observation inferred that apoptosis plays a pivotal role in the death of emodin treated cells at 6 hr, 12 hr, and 24 hr time points compared to untreated cells. However, apoptosis induced after 6 hr is significantly higher than other time points (Fig 10A and 10B)

**Figure 10:**
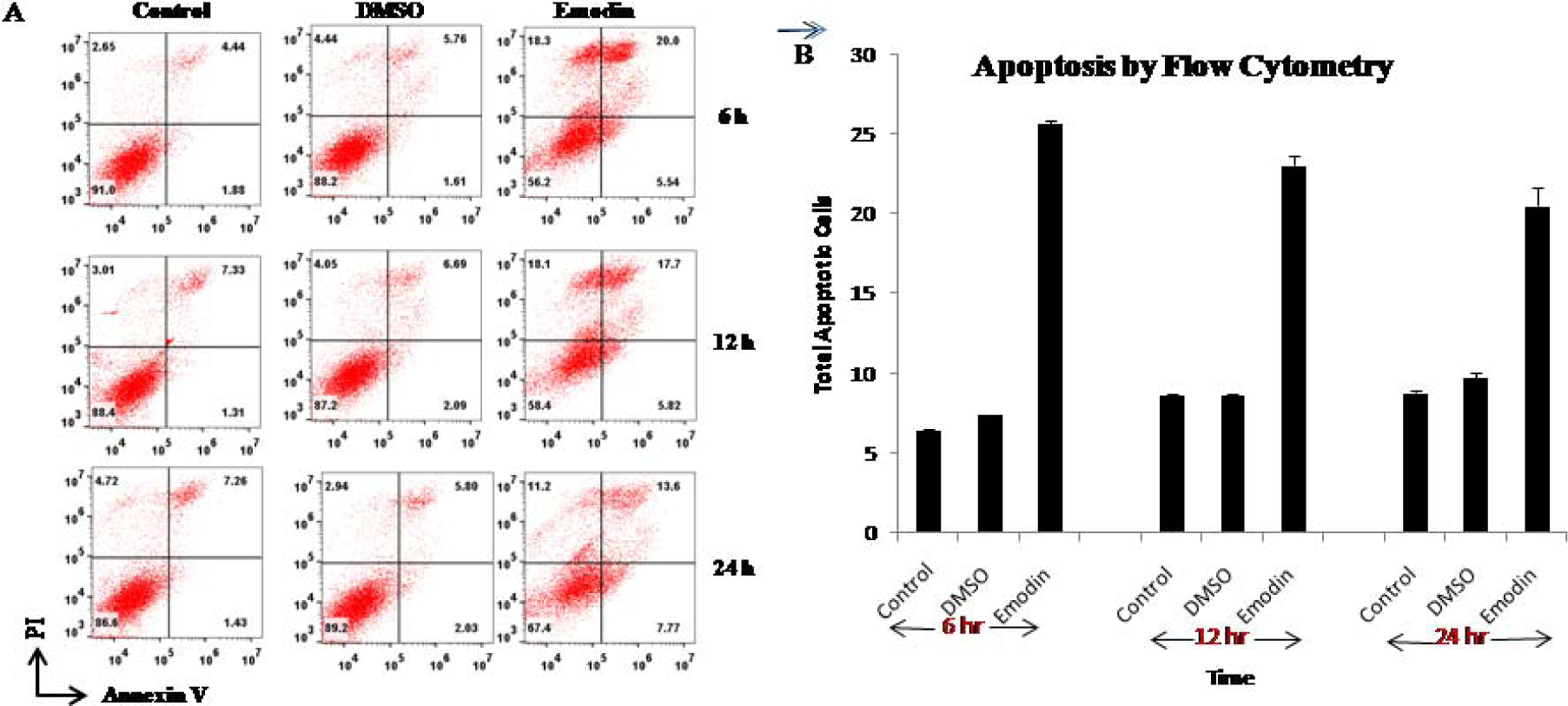
Flow cytometry analysis of emodin treatment on C6 cells. Emodin induces apoptosis at all observed time points. However cell death due to apoptosis is significantly higher after 6hr of emodin treatment.

### 3.7. Emodin treatment induces ROS production

To further substantiate apoptosis induced by emodin in glioma cells, we carried out a ROS detection assay. Overproduction of intracellular ROS causes dysfunction of the mitochondrial membrane and MMP dissipation which ultimately leads to cell death by apoptosis. C6 cells were treated with IC-50 values of emodin for 2 hr, 4 hr, and 6 hr, followed by spectroflourometric analysis. Our results show that emodin induces significant ROS production at all time points compared to control cells (Fig 11).

**Figure 11:**
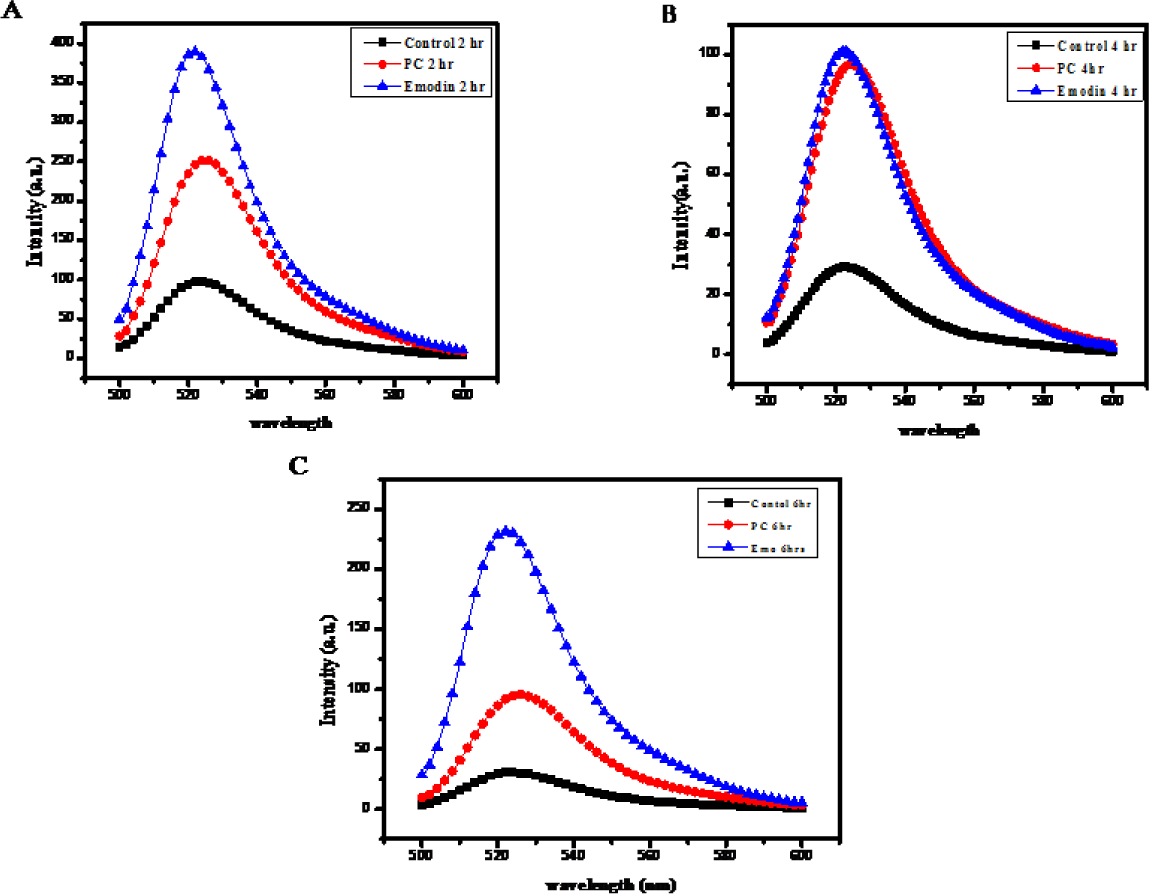
Emodin increases ROS levels: Spectrofluorometer images showing effect of emodin on C6 cells after (A) 2 h (B) 4 h (C) 6 h. There occurs ∼4 fold, ∼3.5 fold and ∼ 8 fold increments in ROS production in C6 cells after 2 h, 4 h and 6 h of 120 μM emodin treatment compared to untreated C6 cells.

### 3.8. Effect of Emodin on EMT in glioblastoma cells

Glioma cells undergoing EMT attain the potential to initiate invasion and metastasis. To evaluate the effect of emodin treatment on EMT, we carried out expression analysis of some important EMT markers at the transcriptional and translational level. For this purpose, total protein and total RNA was isolated after drug treatment of C6 cells at 3 hr, 6 hr, 12 hr and 24 hr and subjected to western blotting and qRT-PCR.

#### 3.8.1. N-Cadherin

N-Cadherin is one of the important EMT markers in glioma and several studies have shown inverse co-relation of glioma invasion with N-cadherin expression. We noticed that after emodin treatment, N-cadherin protein expression levels significantly raise ∼3.5 fold and ∼1.5 fold after 6hr and 12hr time points respectively. Similarly, mRNA expression of N-Cadherin following drug treatment was found to increase by ∼2 fold and ∼ 1.7 fold after 6 hr and 12 hr respectively (Fig. 12 A, B).

**Figure 12:**
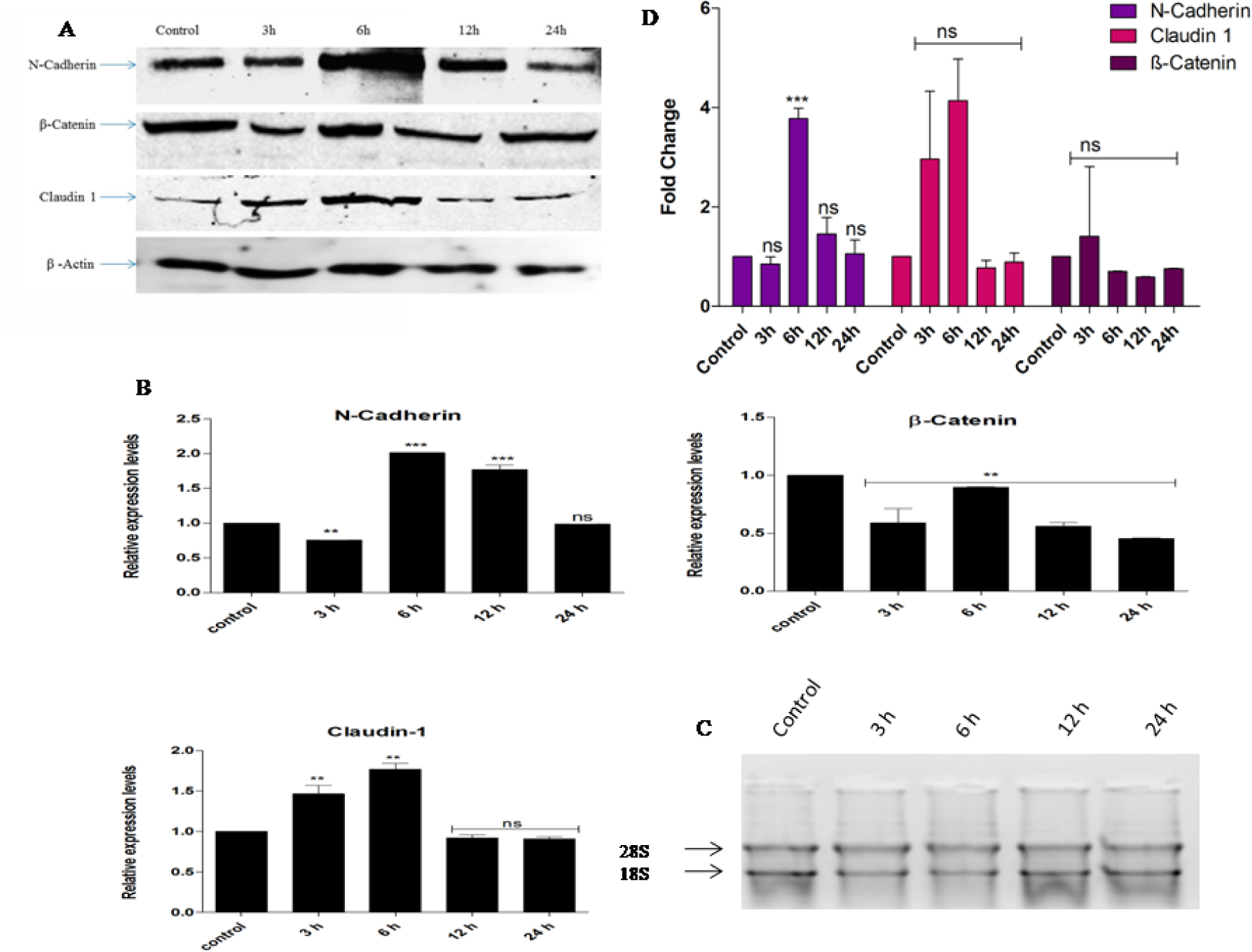
Immuno blots showing the expression of N-cadherin, βcatenin and Claudin-1 proteins in C6 gliom cells after Treatment with Emodin. (A) N-cadherin, βcatenin and Claudin-1 proteins are differentially expressed i c6 cells after emodin treatment.β-actin was used as loading control and expression in each group was normalized to β-actin relative to the normalized expression in control cells. (B) **qRT-PCR for assessing the expression of** N-cadherin, β-catenin and Claudin-1 in Emodin treated C6 glioma cells (C) Gel picture of total RNA isolated from C cells under different conditions. (D) Bar graphs depicting expression levels of N-cadherin, βcatenin and Claudin-proteins as shown by densitometric analysis in arbitrary densitometric units (ADU) in Emodin treated C6 cells. Data are expressed as mean ± SD. Bars indicate standard deviation and asterisk represent significance *P < 0.05; **P< 0.01; ***P< 0.00.1 Dunnett’s Multiple Comparison Test was used to analyze the data.

#### 3.8.2. β-Catenin

β-catenin is a well-established prognostic marker in glioblastoma as its expression levels have been reported to increase both at mRNA and protein levels in glioblastoma and high-grade astrocytomas which correlates with malignancy. Our results showed that emodin treatment decreases β-catenin protein expression at 6hr, 12hr and 24 hr by ∼2 fold, ∼2.5 fold and ∼2 fold respectively, however mRNA expression was found to decrease by ∼ 1.8 fold, ∼1.8 fold and ∼2 fold at 3 hr, 12 hr and 24 hr respectively (Fig. 12 A, B).

#### 2.8.3. Claudin-1

In human glioblastoma, there occurs a decrease in mRNA expression of claudin-1 and a significant reduction of protein level. A reduction in the expression of claudin-1 has been correlated with the aggressiveness of tumors. Our study demonstrates that emodin restores lost claudin-1 protein levels by ∼3fold and ∼4 fold at 3 hr and 6 hr respectively and mRNA expression levels by ∼1.5 fold and ∼ 1.8 fold at corresponding time points (Fig. 12 A, B).

### 3.9. Emodin Inhibits AKT phosphorylation

AKT is the central kinase found to be dysregulated in human cancers. It gets activated mainly by phosphorylation at Thr 308 and Ser 473 which activate two different signaling pathways. However, phosphorylation at Thr 308 is reported to be a more reliable and potential biomarker for its activity in various tumors. AKT is phosphorylated at Thr 308 by PDK1. We analyzed the phosphorylation status of AKT after 15 mins, 30 mins, 1hr, and 2hr of drug treatment. We found that emodin inhibits phosphorylation of AKT at Thr 308 at 1 hr and 2 hr time points as compared to control untreated cells (Fig 13).

**Figure 13:**
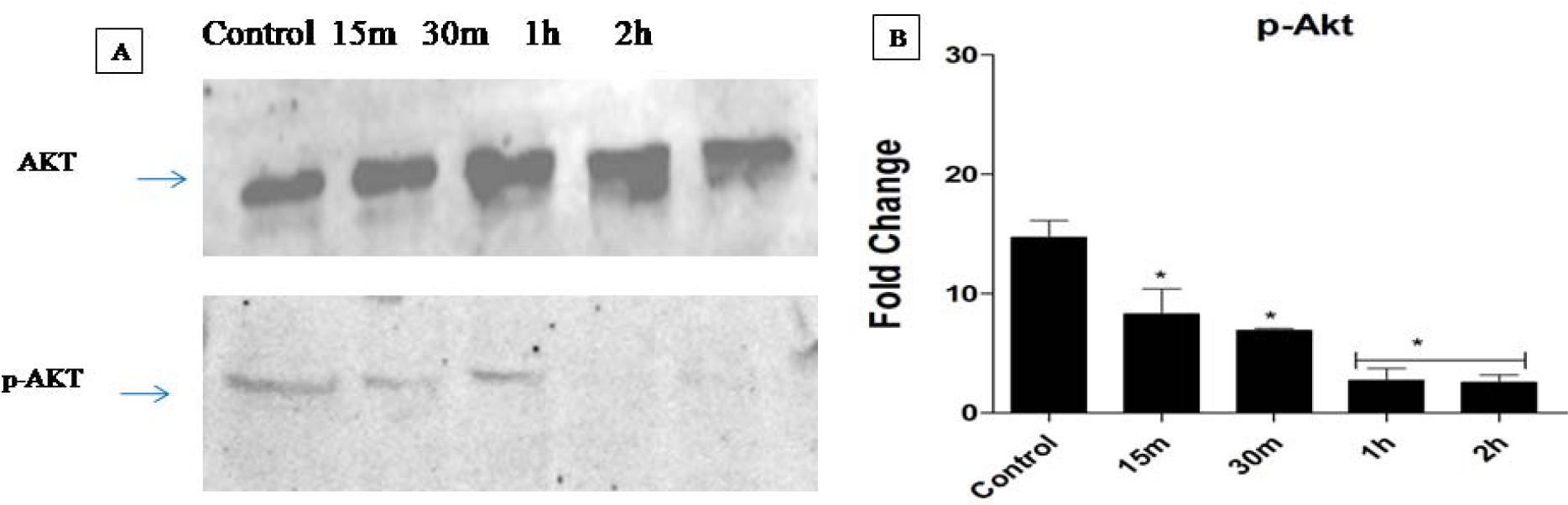
Immuno blots showing the expression of p-AKT protein in C6 glioma cells after Treatment with Emodin. (A) p-AKT protein is downregulated when compared with control. (B) Bar graphs depicting expression levels of p-AKT protein as shown by densitometric analysis in arbitrary densitometric units (ADU) in Emodin treated C6 cells. Data are expressed as mean ± SD. Bars indicate standard deviation and asterisk represent significance*P < 0.05; **P< 0.01; ***P< 0.001. Dunnett’s Multiple Comparison Test was used to analyze the data.

### 3.10. Phospho-Kinase Array evaluation of downstream effectors

Previous results show that emodin inhibits the phosphorylation of Akt at the Thr 308 phosphorylation site after one hour of treatment. To explore potential downstream mediators of different signaling pathways which be affected by emodin treatment, we studied the phosphorylation of 37 kinases (Proteome Profiler Human Phospho Kinase Array) in untreated and emodin treated U87-MG cells at 30 mins, 45 mins, and 60 mins time points. Significant observable upregulation in phosphorylation following emodin treatment was only observed in glycogen synthase kinase 3β (GSK-3 β) (S9) and Lysine deficient protein kinase 1 (WNK1) (T60) (Fig 14).

**Figure 14:**
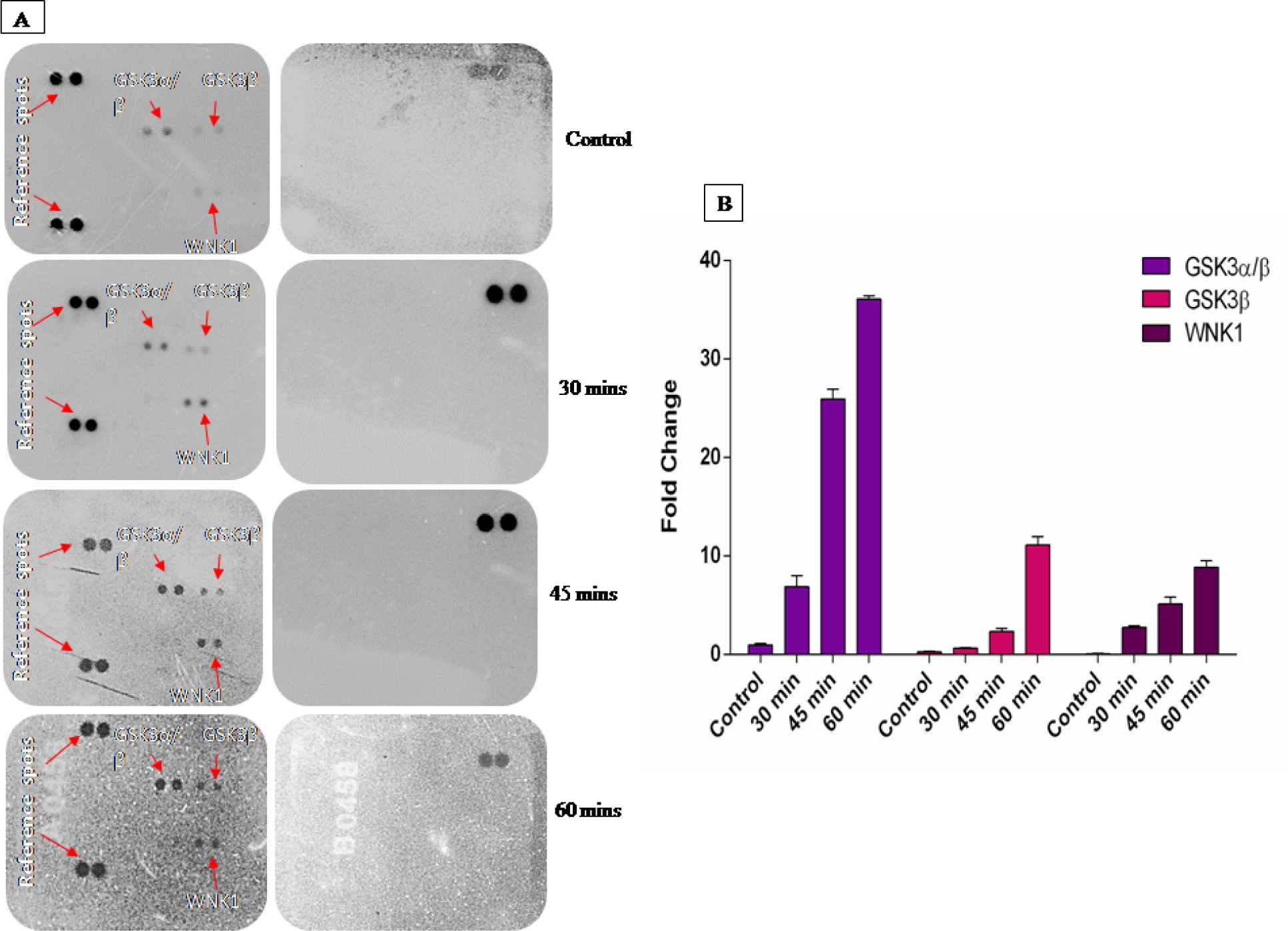
**(A)** A human phospho-kinase blot array of U87MG control and emodin treated cells for 30 mins, 4 mins and 60 mins. The kinases depicted by arrows (red) in the blots represent major signaling protein targets affected by emodin treatment.(B) Representative bar graphs depicting expression levels of GSK3[/β, GSK3βand WNK1 protein as shown by densitometric analysis in arbitrary densitometric units (ADU) in Emodin treated U87-MG cells. Data are expressed as mean ± SD. Bars indicate standard deviation and asterisk represent significance*P < 0.05; **P< 0.01; ***P< 0.001. Bonferroni’s multiple comparisons test was used to analyze the data.

### 3.11. Emodin inhibits nuclear translocation of NF-**ĸ**B

NF-ĸB translocates to the nucleus and activates several genes including genes regulating epithelial to mesenchymal transition in glioblastoma cells. We evaluated the effect of emodin treatment on the nuclear translocation of NF-ĸB after 3hr, 6 hr, 12 hr, and 24 hr. Our immunoblots show that emodin treatment keeps IĸB in active form as indicated by its expression which retains NF-ĸB dimers in the cytoplasm compared to control at all time points (Fig 15). The nuclear expression of NF-ĸB is decreased at all time points which might be the reason for inhibition of EMT at gene expression level.

**Figure 15:**
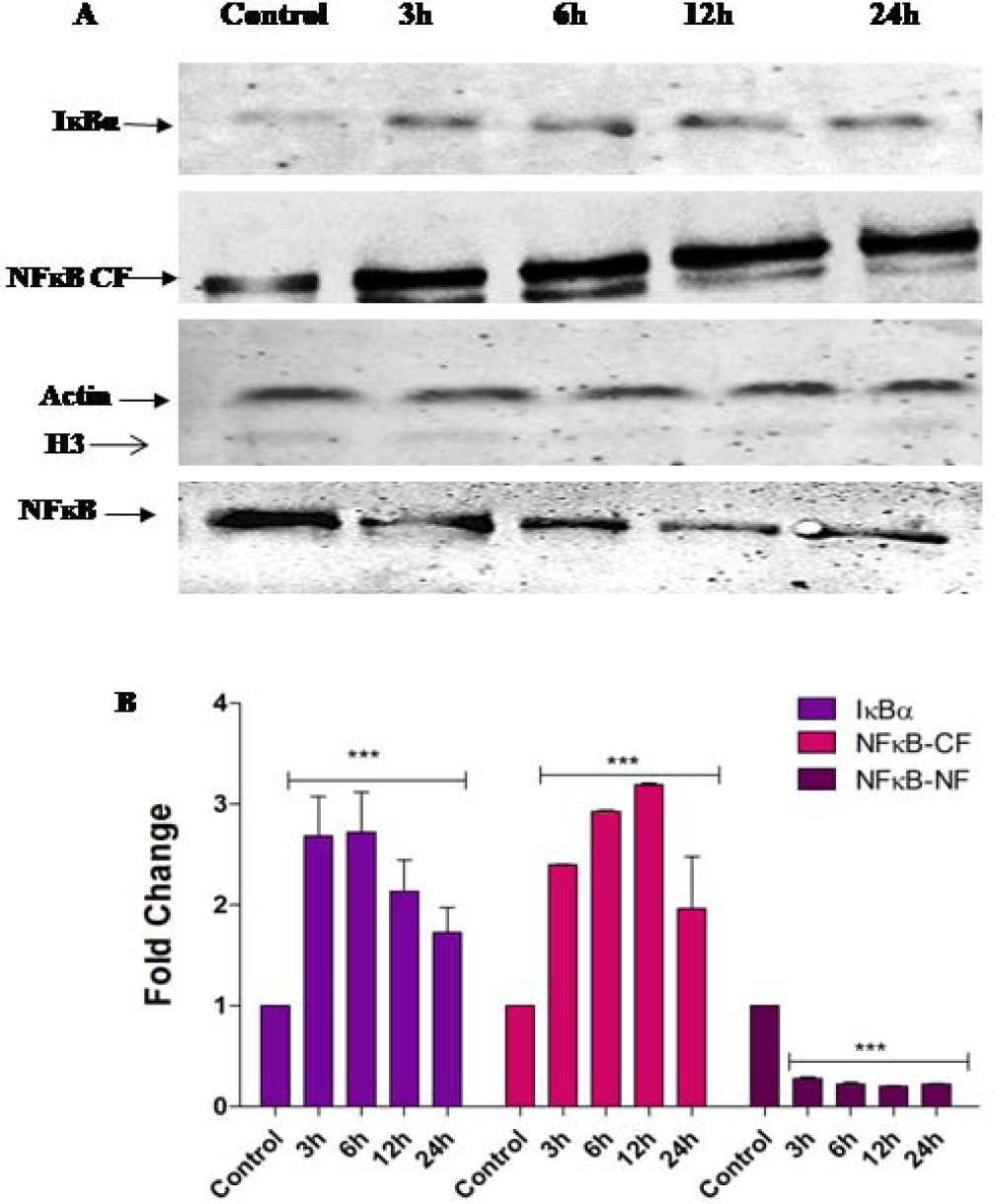
Immuno blots showing the expression of I-ĸB, NF-ĸB protein in C6 glioma cells after Treatment with Emodin. (A) I-ĸB and NF-ĸB (CF) proteins are upregulated when compared with control.β-actin was used as loading control and expression in each group was normalized to β-actin relative to the normalized expression in control cells. NF-ĸB (NF) protein is downregulated when compared with control.H3 was used as loading control and expression in each group was normalized to H3 relative to the normalized expression in control cells.(B) Representative bar graphs depicting expression levels of I-ĸB, NF-ĸB protein as shown by densitometric analysis i arbitrary densitometric units (ADU) in Emodin treated C6 cells. Data are expressed as mean ± SD. Bars indicate standard deviation and asterisk represent significance*P < 0.05; **P< 0.01; ***P< 0.001. Bonferroni’s multiple comparisons test was used to analyze the data.

### 3.12. Molecular Dynamic simulation of PI3K**δ**-Emodin complex

The average RMSD calculated for the protein-ligand complex is normally used to judge the conformational changes and stability pattern of the system in the time scale studies. The stability pattern in terms of RMSD of PI3Kδ protein bound with emodin is shown in Fig 16. The initial RMSD deviation was observed up to 38th ns, and from the remaining nanosecond, the changes in the RMSD were seen. After 39th ns, the major count of RMSD was notified around 0.58 to 0.61 nm. From the RMSD graphs, it was observed that the initial rise in the RMSD may be due to the equilibration phase of the system, and the stability of the protein-ligand complex was obtained from the 39th to 100th ns simulation. In addition, the stability of the ligand in terms of RMSD was also calculated to fetch the binding conformation of emodin inside the binding site of the protein. From the graph, it is observed that the ligand is more comfortable in the active site of protein with the calculated RMSD of 0.05 nm (Fig 17). Further, the fluctuation of amino acid residues in the protein was analyzed. The data suggest that the amino acid residues such as Tyr58, Glu56, Pro57, Leu58, Phe59, His60, Met61, Arg300, Lys400, Ala401, Lys402, Lys403, Ala404, Arg405, Ser406, Thr407, Phe408, Ser411, Lys412 and Lys1028 showed higher fluctuation during the simulation time. The statistical value calculated for RMSF data suggests that the above amino residues are responsible for the protein flexibility and all these residues correspond to the loop regions in the protein (Fig 18). Finally, the stability of the protein-ligand complex in terms of hydrogen bond was calculated and plotted in Figure 18. From the data, it is observed that a maximum of 4 hydrogen bonds are occupied in the protein-ligand complex, in that 2 hydrogen bonds are constant throughout the simulation. Overall, from the analysis, it is confirmed that the emodin was found to be stable inside the binding site of PI3Kδ.

**Figure 16:**
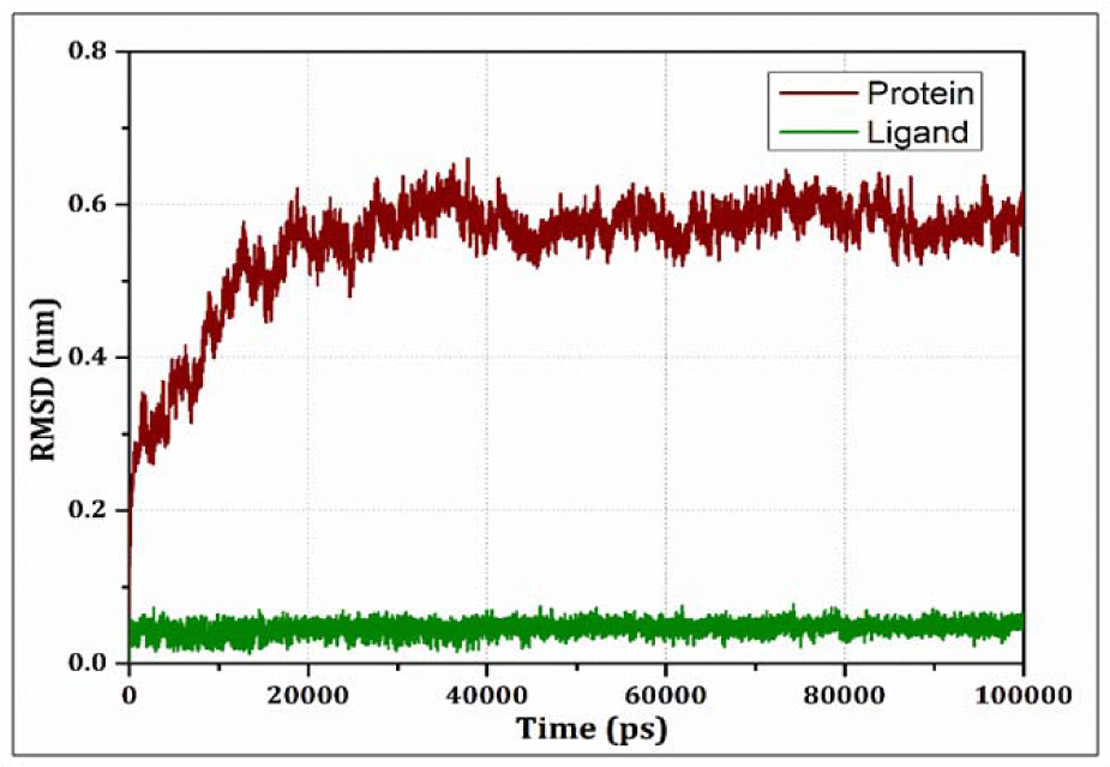
Backbone RMSD calculated for the protein-ligand complex during the simulation time of 100 ns.

**Figure 17:**
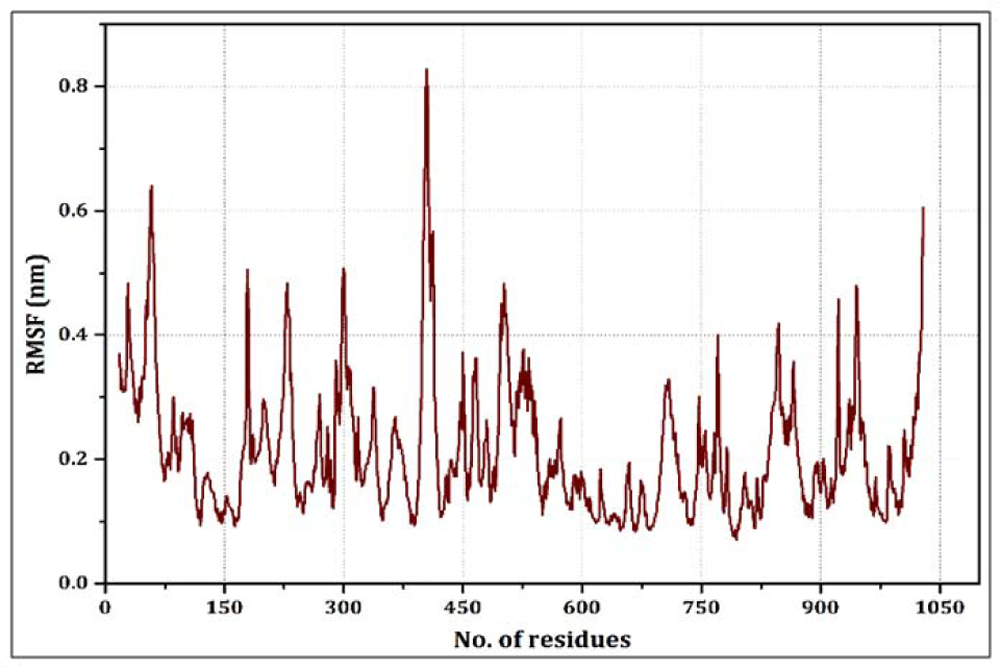
Flexibility of amino acids calculated through RMSF analysis

**Figure 18:**
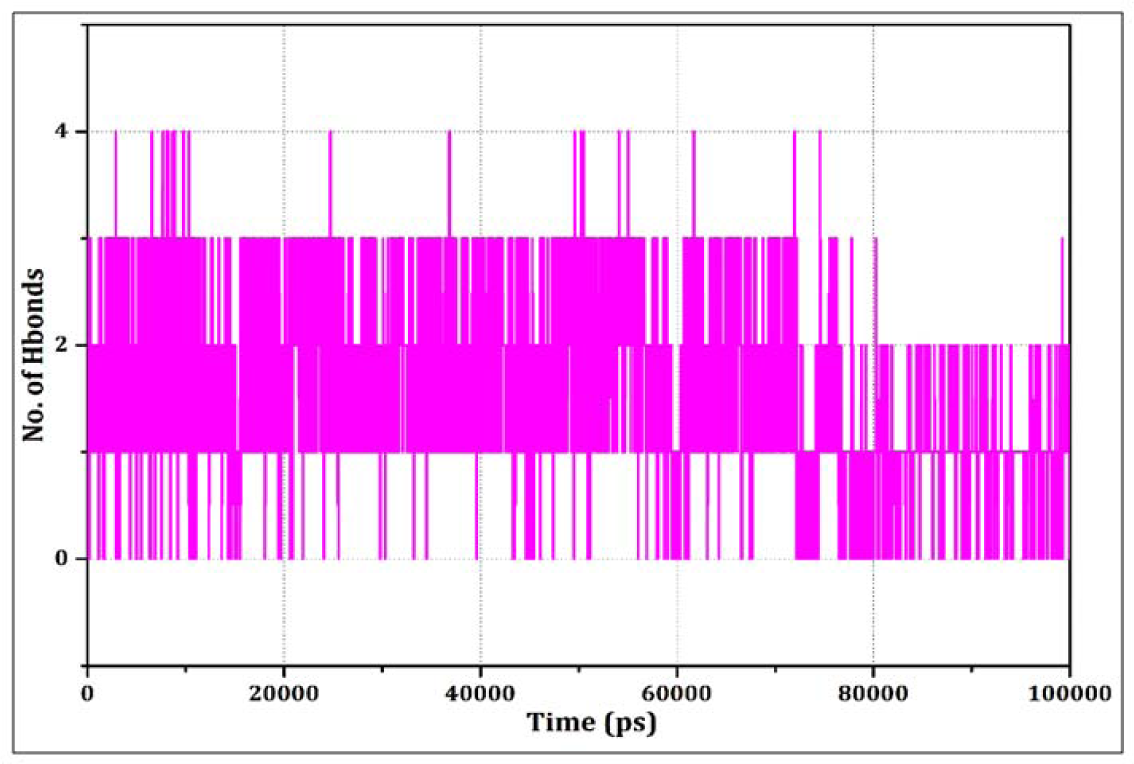
Intra molecular H-bond observed during the protein-ligand complex.

## 4.0 Discussion

Glioma forms a major health challenge worldwide with several signaling transduction pathways being deregulated affecting various cellular processes. Gliomas are often recognized as one of the most heterogeneous and recurrent cancers. They are obstinately resistant to treatment and hence almost incurable. Genetic heterogeneity, and overlapping signal transduction pathways between the heterogenic populations of contributing stromal cells associated with gliomas lead to the limited drug delivery/effect to the tumor and thus counted among several mechanisms underlying therapeutic failure. One of the main pathways deregulated in glioma comprises PI3K-AKT and its associated downstream targets like NF-κB which affects different proteins/transcription factors influencing many aspects of gliomagenesis like epithelial to mesenchymal transition (EMT). Drug development studies to date have revealed only a modest effect in attenuating the growth of these tumors. Ours thus was a combination of *in-silico* and *in-vitro* approach targeted against specific catalytic isoform (p110δ) of Class IA PI3K which seems to be an initial focal point in these deregulated signaling transduction pathways with potent and selective inhibitors.

Initially, we adopted an in-silico approach to screening a range of natural molecules for a potent P110δ inhibitor activity based on sequence similarity of Idelalisib /Cal-101, which is a synthetic P110δ inhibitor approved for use. We carried out protein preparation, ligand preparation, molecular docking, calculation of binding free energy, and ligand interaction profile generation. We shortlisted three hits based on a higher docking score and delta G binding energy compared to our positive control. Among these three hits, Emodin, a naturally occurring anthraquinone derivative predominantly abundant in the roots of Rheum *Palmtum*, Polygonum *cuspidatum*, and Polygonum *multiflorum* were found to exert a strong antitumor effect on glioblastoma cells. Treating glioma cells with IC-50 values of emodin exert a strong antitumor effect in glioma cells. It inhibits cell proliferation and induces cell death as observed in human esophageal cancer cells [44]. Similarly, cell proliferation inhibition as well as cell death-inducing potential was observed in U87-MG and C6 glioma cell lines. Emodin has been found to induce apoptosis in human cervical cancer cells and other tumors [45]. Accordingly, we explored the role of emodin in inducing apoptosis in glioma cells. We carried out propidium iodide staining which demonstrated that emodin induces morphological changes typical of apoptosis and nuclei stained with PI in emodin-treated cells were far more as compared to untreated cells. Furthermore, Annexin V FITC flow cytometry analysis confirms that emodin treatment causes cell death by apoptosis. Emodin has been found to generate ROS in gastric carcinoma cells [46]. To understand the mode of apoptosis induction, we evaluated ROS production in glioma cells. We observed that emodin significantly induces ROS production in glioma cells which may be an emodin-mediated apoptosis-inducing factor in glioma cells.

GSK-3β activity is implicated in drug resistance in various cancers. It is inactivated by phosphorylation at S9 by various kinases such as PKA, Akt, and p70S6K [47]. Inhibition of GSK-3β activity attenuates proliferation, enhanced tumor cell apoptosis, inhibited cell survival, reduced cell motility and clonogenicity in GBM [48]. Our results indicate that emodin treatment upregulates the S9 phosphorylated form of GSK-3β which results in inhibition/ degradation of downstream proteins and transcription factors associated with the growth and proliferation of glioma cells. AKT forms a central signaling hub that regulates more than 100 downstream target proteins, thereby affecting cell growth, survival, cell metabolism, and proliferation. Unregulated overexpression of phosphorylated AKT (p-AKT) is a widespread feature in early as well as advanced cancers [49]. An important transcription factor involved in inducing the EMT process regulated by the activation of Akt is NF-κB. It remains in an inactive state in the cytoplasm bound by Inhibitor kappa B (IkB) proteins such as Ik-Bα, thus inhibiting the EMT process [50]. Our results indicate that emodin deactivates phosphorylated AKT, and inhibits nuclear translocation of NF-κB by keeping it in a bound state with Ik-Bα in the cytoplasm. This inactivation and inhibition of prominent proteins are assumed to have an impact on various protumorigenic processes including EMT. Therefore, we evaluated the expression changes of some important EMT markers. N-Cadherin is one of the important EMT markers in glioma and several studies have shown inverse corelation of glioma invasion with N-cadherin expression [51]. Our results indicate that emodin treatment upregulates N-cadherin expression both at mRNA and protein levels after 6hr and 12 hr treatment. β-catenin shows membranous expression in normal glial cells, however, in tumorous cells its expression is mostly nuclear/cytoplasmic. The unusual expression of β-catenin in cytoplasm/ nucleus could be considered as an indicator of its aberrant expression which indicates an activated Wnt/ β-catenin signaling pathway [52]. Treatment of glioma cells with emodin shows decreased cytoplasmic expression of β-catenin after 12hr compared to control both at mRNA as well as protein levels. Claudin-1 is an important epithelial marker and one of the most important blood-brain barrier components. In tumor cells, this barrier is altogether disturbed allowing unaccounted movement of solutes inside the cells [53]. However, emodin treatment has helped to restore the lost expression of this crucial protein. Treating cells with emodin results in changed expression profiles of these EMT markers which probably counter relevant processes involved in promoting gliomagenesis.

## Conclusion

The current study was focussed at looking for a therapeutic agent that may reduce tumor characteristics of glioma in-vitro. Taken in totality, our results suggest that emodin therapy remarkably reduces the proliferation, invasion and induces apoptosis in glioma cells possibly through different mechanisms, targeting multiple pathways involved in tumor growth, and development. Our results provide a basis for further evaluation of emodin on animal models of glioma, and thus may pave way for its use in clinical trials later on.

## Acknowledgements

The authors thank Umar Majeed Wani and Mir Khurshid Iqbal for their suggestions and technical assistance. The authors also thank Umer Mehraj for helping with FACS analysis.

## Funding

The work was supported by Department of Biotechnology, Government of India.

EMT: epithelial to mesenchymal transition
CBTRUS: Central Brain Tumor Registry of United States
TMZ: Temozolomide
NCCN: National Comprehensive Cancer Network
GBM: Glioblastoma Multiforme
Iĸ-B: Inhibitors of NF-ĸB
MMP: Matrix Metalloproteinase
MM-GBSA: Prime/molecular Mechanics Generalized Born-Surface Area
GROMACS: Groningen Machine for Chemical Simulation
NVT: Isothermal-Isochoric
NPT: Isothermal-Isobaric
PME: Particle Mesh Ewald
RMSD: Root Mean Square Deviation
RMSF: Root Mean Square Fluctuation
EMEM: Minimum Essential Medium Eagle
FBS: Fetal Bovine Serum
MTT: 3-(4,5-dimethylthiazol-2-yl)-2,5-diphenyltetrazolium bromide
DMSO: Dimethyl Sulphoxide
LDH: Lactate dehydrogenase
GSK-3 β): Glycogen Synthase Kinase 3β
WNK1: Lysine Deficient Protein Kinase-1

## Notes

### Competing Interest Statement

The authors have declared no competing interest.

